# The Ty1 integrase nuclear localization signal is necessary and sufficient for retrotransposon targeting to tRNA genes

**DOI:** 10.1101/2019.12.18.879569

**Authors:** Amna Asif-Laidin, Christine Conesa, Amandine Bonnet, Camille Grison, Indranil Adhya, Rachid Menouni, Hélène Fayol, Noé Palmic, Joël Acker, Pascale Lesage

## Abstract

Integration of transposable elements into the genome is mutagenic. Mechanisms that target integration into relatively safe locations and minimize deleterious consequences for cell fitness have emerged during evolution. In budding yeast, the integration of the Ty1 LTR retrotransposon upstream of RNA polymerase III (Pol III)-transcribed genes requires the interaction between the AC40 subunit of Pol III and Ty1 integrase (IN1). Here we show that the IN1-AC40 interaction involves a short linker sequence in the bipartite nuclear localization signal (bNLS) of IN1. Mutations in this sequence do not impact the frequency of Ty1 retromobility, instead they decrease the recruitment of IN1 to Pol III-transcribed genes and the subsequent integration of Ty1 at these loci. The replacement of Ty5 retrotransposon targeting sequence by the IN1 bNLS induces Ty5 integration into Pol III-transcribed genes. Therefore, the IN1 bNLS is both necessary and sufficient to confer integration site specificity on Ty1 and Ty5 retrotransposons.

## INTRODUCTION

Transposable elements (TEs) are mobile repetitive DNA sequences found in the genomes of most organisms (Huang et al., 2013). TEs are mutagenic and represent a threat to genome integrity, inactivating or altering host gene expression or inducing large chromosomal rearrangements (Bourque, 2009; Levin and Moran, 2011). In humans, more than hundred heritable diseases have been assigned to *de novo* TE insertions (Hancks and Kazazian 2016). TEs also play a role in genome evolution by modifying host functions, phenotypes, and gene regulation, and can contribute to the long-term adaptation of organisms to different environments (Chuong et al., 2016).

Where TEs integrate in the genome will determine their impact on their host. TE distribution, which is rarely random (Sultana et al., 2017), arises from the balance between two processes. First, selection leads to the elimination of strongly deleterious insertions, and the maintenance of beneficial ones (Chuong et al., 2016; Cosby et al., 2019). Second, TEs have repeatedly evolved mechanisms that direct their integration into “safe” locations, where insertions will have minimal adverse effect on the organism’s fitness (Boeke and Devine, 1998; Cheung et al., 2018; Spaller et al., 2016). These regions often consist of non-essential repeated sequences, such as telomeric regions, ribosomal DNA arrays, and transfer RNA genes (*tDNA*s), or non-essential regions upstream of open reading frames (Baller et al., 2012; Fujiwara et al., 2005; Guo and Levin, 2010; Kling et al., 2018; Mularoni et al., 2012; Naito et al., 2009; Pardue and DeBaryshe, 2011; Penton and Crease, 2004; Ye et al., 2005; Zou et al., 1996). Preferential targets have been described for the integration of different classes of TEs, including retroelements (Sultana et al., 2017).

Long terminal repeat (LTR) retrotransposons are retroelements related to retroviruses. They replicate by reverse transcription of their mRNA into a double-stranded DNA copy (cDNA), which is imported into the nucleus and integrated into the genome by the LTR-retrotransposon integrase (IN). The interaction between IN and cellular tethering factors plays a major role in integration site selection by targeting pre-integration complexes (PICs) to specific chromosome locations. Tethering factors were first identified for Ty3 and Ty5 in *S. cerevisiae*. These LTR retrotransposons integrate at the transcription start site of Pol III-transcribed genes and in subtelomeric regions, respectively (Kirchner et al., 1995; Xie et al., 2001). The Tf1 LTR retrotransposon of *Schyzosaccharomyces pombe* and the MLV retrovirus both target the promoter region of Pol II-transcribed genes (Gupta et al., 2013; Hickey et al., 2015; Jacobs et al., 2015; De Rijck et al., 2013; Sharma et al., 2013), and the HIV-1 retrovirus targets the gene body of Pol II-transcribed genes (Cherepanov et al., 2003; Llano et al., 2006). In all cases, tethering factors bind chromatin or have functions related to DNA transcription or replication (Sultana et al., 2017).

Ty1, the most active and abundant LTR-retrotransposon in *S. cerevisiae*, integrates preferentially within a 1kb window upstream of Pol III-transcribed genes. It targets nucleosomal DNA near the H2A/H2B interface (Baller et al., 2012; Mularoni et al., 2012). This integration pattern allows Ty1 to replicate while minimizing disruption to the host genome, as most Pol III-transcribed genes are multicopy *tDNA*s and thus individually non-essential. Furthermore, Ty1 insertion has a limiting impact on *tDNA* expression (Bolton and Boeke, 2003). Targeted integration proximal to *tDNA*s is a strategy that has been adopted several times by TEs to minimize damage to compact genomes (Cheung et al., 2018; Kling et al., 2018). The integration of Ty1 in these regions requires a functional Pol III promoter in the target gene (Devine and Boeke, 1996) and is influenced by the chromatin-remodeling factor Isw2 and the Bdp1 subunit of TFIIIB (Bachman et al., 2005).

Recently, we have shown that an interaction between Ty1 IN (IN1) and the AC40 subunit of Pol III is a major driver for Ty1 integration upstream of Pol III-transcribed genes (Bridier-Nahmias et al., 2015). The study used the *S. pombe* AC40 ortholog (AC40sp) as a loss-of-interaction mutant. The replacement of AC40 by AC40sp severely compromised Ty1 integration upstream of Pol III-transcribed genes, leading to a redistribution of Ty1 insertions in the genome. IN1 binding to other Pol III subunits was also described *in vitro* (Cheung et al., 2016). However, it is not clear whether these interactions participate to Ty1 integration site selection.

IN1 has a three domains organization common to all retroelement integrases; the Zn^2+^ coordinating N-terminal domain (NTD), the catalytic core domain (CCD) and the less conserved C-terminal domain (CTD) (Figure 1A, top) (Wilhelm et al., 2005). IN1 C-terminal residues 578-635 are necessary and sufficient to mediate the interaction with AC40 *in vivo* (Bridier-Nahmias et al., 2015). This region also contains a bipartite nuclear localization signal (bNLS, residues 596-630; Figure 1A, top) consisting of two Lys-Lys-Arg motifs separated by a 29 amino-acid linker (Kenna et al., 1998; Lange et al., 2011; Moore et al., 1998). This raises the question of whether IN1 nuclear import and interaction with AC40 could act in concert during Ty1 replication.

**Figure 1.**
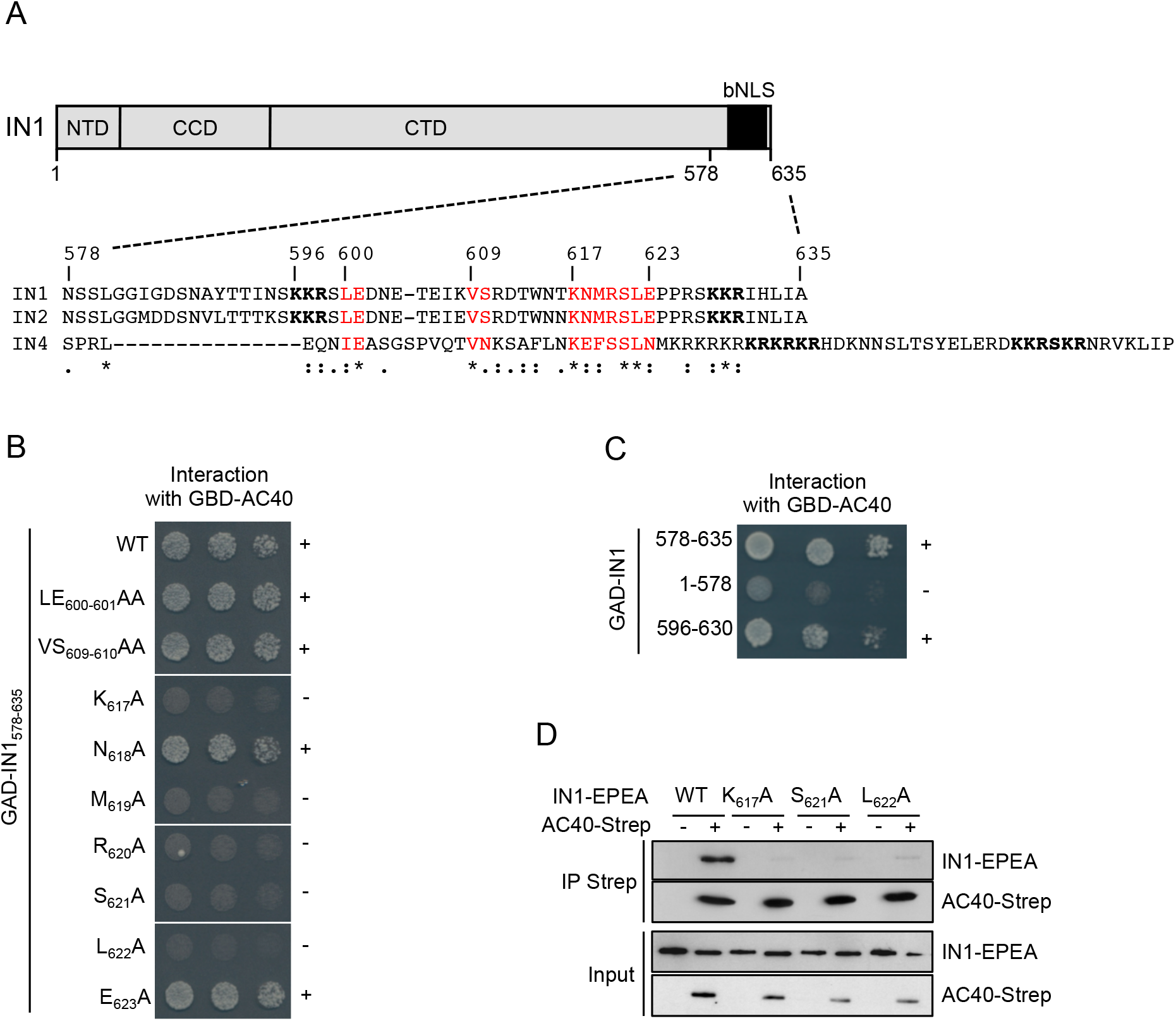
bNLS linker sequence mutations abolish the interaction with AC40. **A.** *Top*. The Ty1 integrase (IN1) showing N-terminal and catalytic core domains (NTD and CCD) and the bipartite NLS at the C-terminus (CTD). *Bottom*. Alignment of amino acid sequences of Ty1, Ty2, and Ty4 integrase C-termini (IN1, IN2, and IN4, respectively). In bold: basic amino acids required for NLS function. In red: amino acids in the NLS linker relatively conserved between the three integrases. *, Identity; :, high similarity; ., low similarity; -, gap in sequence. **B.** Two-hybrid interaction between GBD-AC40 and WT or mutant GAD-IN1_578-635_. Alanine substitutions in IN1_578-635_ are indicated. Cells were plated in two-fold serial dilutions on DO-Leu-Trp-His plates to detect interaction. No growth or protein expression defects were detected (Figure S1A-B). +, Interaction; -, no interaction. **C.** Two-hybrid interaction between GBD-AC40 and different IN1 regions fused to GAD, as indicated. Cells were plated in ten-fold serial dilutions on DO-Leu-Trp-His plates to detect interaction. No growth or protein expression defects were detected (Figure S1C-D). +, Interaction; -, no interaction. **D.** In vitro interaction between AC40 and IN1 proteins co-expressed in *E. coli*. Immuno-precipitation of protein extracts was performed from bacteria cells expressing IN1-EPEA or K_617_A, S_621_A, or L_622_A IN1-EPEA mutants alone (-) or together with AC40-Twin-Strep-tag (+). Cell lysates (Input) and immunoprecipitates (IP Strep) were analyzed by Western blot with anti-Strep-Tactin antibody and CaptureSelect™ Biotin Anti-C-tag Expected sizes are 41 kDa for AC40-Twin-Strep-tag and 100 kDa for IN1-EPEA (WT and mutants).

In this study, we identify a short sequence in the bNLS linker of IN1 that directs the interaction with AC40. Single amino acid substitutions in this sequence do not affect the frequency of Ty1 retrotransposition but impair the recruitment of IN1 to Pol III-transcribed genes. Consequently, these IN1 mutations induce the same changes in the Ty1 integration profile as observed in the AC40sp loss-of-interaction mutant. When the IN1 bNLS is used to replace the Ty5 IN sequence responsible for Ty5 integration into heterochromatin, Ty5 integration is re-directed to Pol III-transcribed genes. This work therefore confirms the fundamental role of the IN1-AC40 interaction in Ty1 integration site selection and reveals that the IN bNLS is necessary and sufficient to confer Ty1 integration preference to another retrotransposon.

## RESULTS

### bNLS linker sequence mutations abolish the interaction with AC40

*S. cerevisiae* LTR-retrotransposons Ty1, Ty2, and Ty4 have the same integration preferences for regions upstream of Pol III-transcribed genes (Carr et al., 2012; Kim et al., 1998) and the C-termini of their integrases (IN1, IN2, and IN4, respectively) interact with the Pol III subunit AC40 (Bridier-Nahmias et al., 2015). To identify conserved amino acids potentially involved in the AC40 interaction, we aligned the C-terminal sequences of IN1, IN2, and IN4 (Figure 1A, bottom) and observed that IN1 and IN2 are highly similar in this region, whereas IN4 is more divergent. Amino acids at positions 600-601, 609-610, and 617-623 in IN1 were either identical or highly similar in all three INs. We replaced each of these amino acids by alanine, individually or in pairs, in a <u>G</u>al4 <u>a</u>ctivating <u>d</u>omain GAD-IN1_578-635_ fusion protein and studied the interaction of the mutant fusion proteins with <u>G</u>al4 <u>b</u>inding <u>d</u>omain GBD-AC40 using a two-hybrid assay. The interaction between IN1_578-635_ and AC40 was maintained in the presence of mutations LE_600-601_AA, VS_609-610_AA, N_618_A or E_623_A, and suppressed by single alanine substitution of K_617_, M_619_, R_620_, S_621_ or L_622_ (Figure 1B). GBD-AC40 interacted with GAD-IN1_578-635_ but not with GAD-IN1_1-578_, as shown previously (Bridier-Nahmias et al., 2015). Since amino acids K_617_-L_622_ are located in the IN1 bNLS linker sequence, we also tested the interaction between GBD-AC40 and GAD fused to the entire bNLS sequence (GAD-IN_596-630_). Interaction between the two fusion proteins was detected, suggesting that this region of 34 amino acids in IN1 is required for interaction with AC40 (Figure 1C).

To determine whether amino acids required for the two-hybrid interaction between AC40 and the IN1 C-terminus were also critical for the interaction between the two full-length proteins, we co-expressed AC40-strep and WT or mutant IN1-EPEA (Glu-Pro-Glu-Ala) tagged proteins in *E. coli* and performed immunoprecipitation of the purified proteins. We focused on K_617_, S_621_ and L_622_, given their strict conserved in IN1, IN2 and IN4 (Figure 1A, bottom). In the presence of Strep-Tactin beads, AC40-strep co-immunoprecipitated with WT IN1-EPEA but not with the IN1-EPEA K_617_A, S_621_A, and L_622_A mutants (Figure 1D). Thus, full-length IN1 and AC40 proteins bind directly to each other, and their interaction depends on residues K_617_, S_621_ and L_622_ located in the IN1 bNLS.

### Non-AC40 binding IN1 mutants do not affect Ty1 integration frequency

Mutations in the IN1 bNLS that induce a substantial or complete loss of IN1 nuclear accumulation reduce the frequency of Ty1 retromobility, as seen when the two Lys-Lys-Arg (KKR) motifs are mutated, either individually or simultaneously, but also for mutations of specific acidic residues in the linker region (Kenna et al., 1998; Lange et al., 2011; Moore et al., 1998). To investigate whether the bNLS amino acids we identified as being necessary for the AC40 interaction were also required for Ty1 nuclear import and retromobility, we introduced K_617_A, S_621_A or L_622_A mutations into a GFP_2_-bNLS fusion protein previously used to assess IN1 bNLS function (McLane et al., 2008). In contrast to the IN1-bNLSmut construct (_596_KKR_598_-AAA and _628_KKR_630_-AAA), the three IN1-bNLS single mutants were still able to target GFP_2_ into the nucleus (Figure 2A). Thus, amino acids required for the IN1-AC40 interaction are dispensable for NLS function.

**Figure 2.**
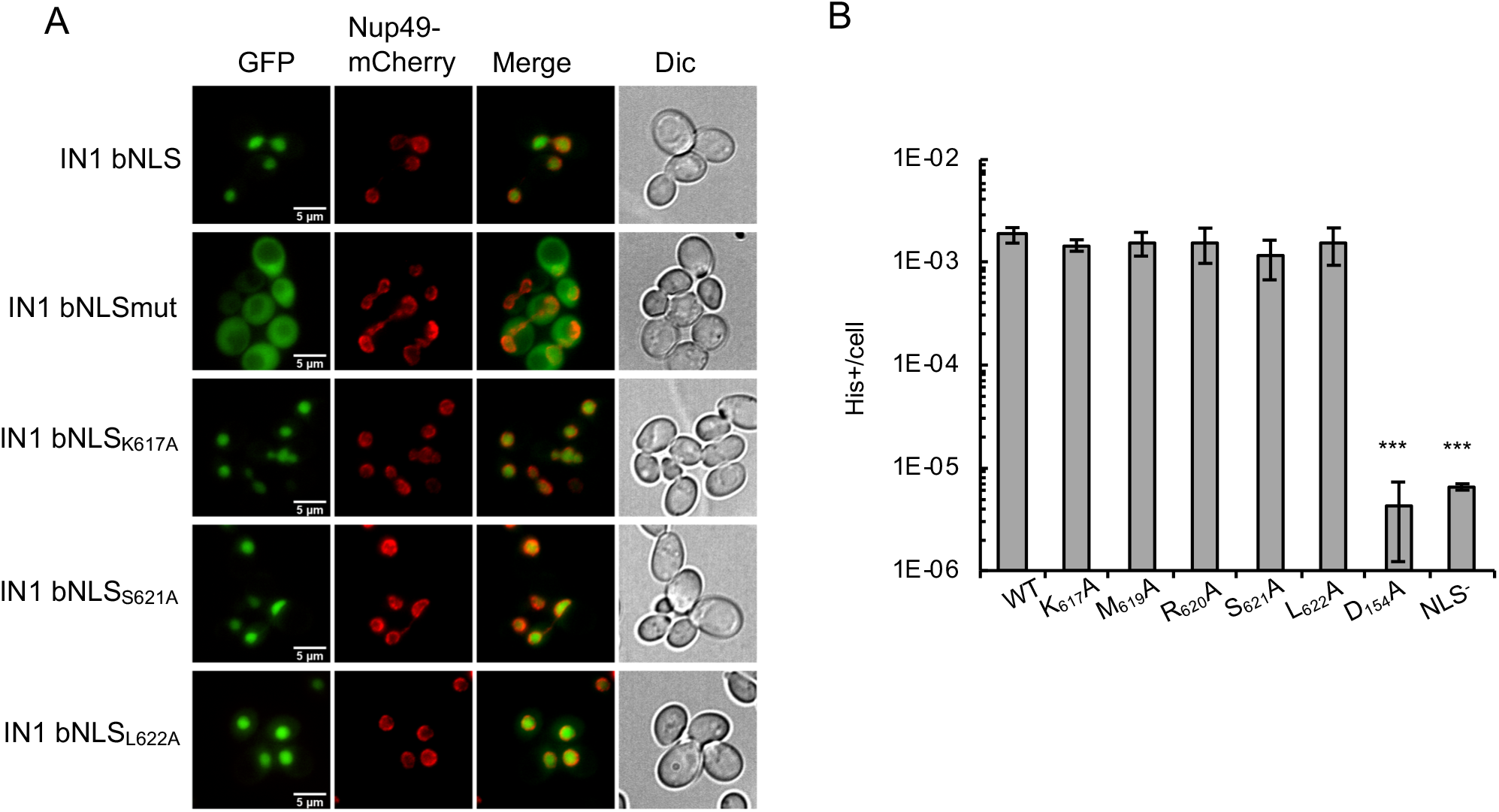
Non-AC40 binding IN1 mutants do not affect Ty1 integration frequency. **A.** Localization of GFP_2_-IN1 bNLS variants. Yeast cells expressing GFP_2_-IN1 bNLS variants and Nup49-mCherry were analyzed by direct fluorescence microscopy. The C-terminal 54 amino acids of IN1 containing the bNLS were fused to the C-terminus of two tandem GFPs (GFP_2_-IN1 bNLS) to create a reporter protein that would be too large for passive diffusion through NPCs. The GFP2-IN1 bNLS_mut_ construct is mutated for both key regions of the bNLS (_596_KKR_598_-AAA and _628_KKR_630_-AAA). Nup49-mCherry signal was used to visualize the location of the nuclear envelope. Corresponding DIC images are shown. **B.** Retrotransposition frequency (log scale) of p*GAL1*-Ty1*his3AI* bearing substitutions of conserved residues by alanine (K_617_A, M_619_A, R_620_A, S_621_A, and L_622_A) in a *spt3-101 rad52∆* strain. IN1 catalytic core domain mutant D_154_A is defective for integration and NLS mutant (KKR_628-630_GGT) for nuclear import. Values are an average of four experiments, each performed with four independent colonies. *p ≤ 0.05, **p ≤ 0.01, and ***p ≤ 0.001; Student t-test.

The same mutations were introduced individually into a Ty1 element containing the retromobility indicator gene *his3AI*, allowing detection of Ty1-*HIS3* insertion events as His^+^ prototroph cells and expressed from the *GAL1* promoter in a 2-micron plasmid (Curcio and Garfinkel, 1991). To determine the frequency of Ty1*his3AI* integration in the genome, we expressed this plasmid in a *spt3-101* null *rad52∆* mutant strain, deficient in endogenous Ty1 expression and homologous recombination. *SPT3* is required for Ty1 transcription and its absence prevents the trans-complementation of the mutant IN1 by WT IN1 from endogenous Ty1 elements (Winston et al., 1984). *RAD52* deletion precludes insertion of the Ty1-*HIS3* cDNA by homologous recombination with preexisting genomic Ty1 copies, a preferred pathway when IN1-dependent integration is defective (Sharon et al., 1994). The frequency of His^+^ cells was similar between strains expressing WT or K_617_A, M_619_A, R_620_A, S_621_A or L_622_A mutant Ty1*his3AI* (Figure 2B). In contrast, mutations that inactivate IN1 nuclear import (Moore et al., 1998) or catalytic activity (Wilhelm and Wilhelm, 2005) caused a substantial decrease in the frequency of His^+^ cells compared to WT (Figure 2B, mutants KKR_628-630_GGT and D_154_A, respectively).

Thus, single amino acid mutations in the linker of the IN1 bNLS that prevent the interaction with AC40 do not impair Ty1 retrotransposition. These data confirm that the IN1-AC40 interaction is not required for Ty1 overall integration frequency (Bridier-Nahmias et al., 2015).

### AC40 recruits IN1 at Pol I and Pol III-transcribed genes

In addition to AC40, other Pol III subunits have been suggested to mediate the interaction between IN1 and Pol III (Cheung et al 2016). To determine whether IN1 remains associated with Pol III in the absence of the IN1-AC40 interaction, we immunoprecipitated Pol III (Oficjalska-Pham et al., 2006) in yeast cells expressing hemagglutinin (HA)-tagged C160, the largest Pol III subunit, and WT or mutant IN1 fused to streptavidin (IN1-strep). WT IN1 was associated to Pol III but not the K_617_A, S_621_A or L_622_A IN1 mutants (Figure 3A). Therefore, the interaction with AC40 is necessary for IN1 binding to Pol III *in vivo*.

**Figure 3.**
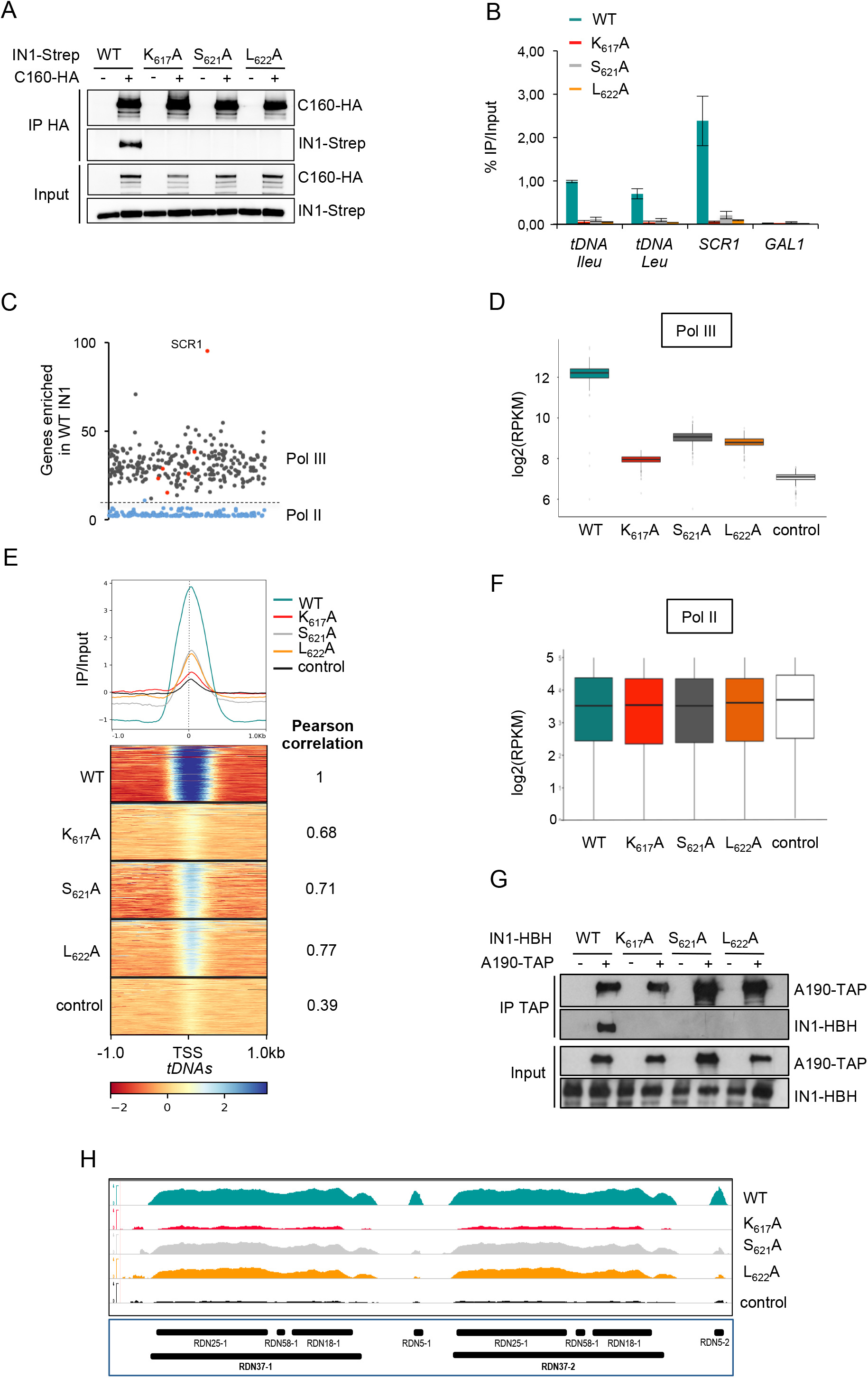
AC40 recruits IN1 at Pol I and Pol III-transcribed genes. **A.** Co-immunoprecipitation between endogenous Pol III (C160-HA) and ectopic IN1 tagged with streptavidin (IN1-Strep) expressed from the *GAL1* promoter. Protein extracts of yeast cells expressing HA-tagged (+) or untagged (-) C160 respectively, and WT or K_617_A, S_621_A, or L_622_A IN1 mutants were incubated with anti-HA coated beads. Cell lysates (Input) and immunoprecipitates (IP HA) were analysed by immunoblotting with anti-HA and anti-Strep antibodies. Expected sizes are 160kDA for C160-HA and 100kDa for IN1-strep (WT and mutants). **B.** Quantitative ChIP analysis of HA-IN1 enrichment at Pol III-transcribed genes. Immunoprecipitated DNA from yeast cells producing ectopic IN1 is expressed as a value relative to that of the input. Pol III transcribed-genes: *tDNA*-Leu and *tDNA*-Ile families (22 and 16 genes, respectively) and the unique *SCR1* gene. *GAL1* ORF serves as a control. The mean values and standard deviation (indicated by error bars) of at least three independent experiments are shown. **C.** Fold enrichment of WT HA-IN1 over input at all the genes where HA-IN1 has been detected by ChIP-seq analysis. Each dot represents a gene. Grey, *tDNA*s; Red, other Pol III-transcribed genes; Blue, Pol II-transcribed genes. **D.** WT and mutant IN1 association with all Pol III-transcribed genes (tRNA and ncRNA genes). Values obtained from ChIP seq analysis have been normalized in log2 RPKM (reads per kilobase per million mapped reads). Control is anti-HA immunoprecipitation of chromatin extracts in cells expressing IN1-Strep. **E.** Genome-wide occupancy profiles (top) and heatmaps (bottom) of WT and HA-IN1 mutants over a 1kb window upstream and downstream of all *tDNA*s. Average ChIP-seq signals have been computed per 10bp bin, normalized to input and adjusted in log2 RPKM. Pearson correlation values corresponding to Figure S2B are indicated. **F.** WT and mutant IN1 association to all Pol-II transcribed genes respectively. Values obtained from ChIP seq analysis have been normalized in log2 RPKM. Control, as described for panel D. **G.** Co-immunoprecipitation between endogenous RNA Pol I (TAP-tagged-A190) and ectopic Ty1 integrase (IN1-HBH). Protein extracts of yeast cells expressing TAP-tagged (+) or untagged (-) A190 and transformed with pCM185 derivatives expressing WT or K_617_A, S_621_A, or L_622_A IN1-HBH mutants were incubated with IgG beads. Cell lysates (Input) and immunoprecipitates (IP TAP) were revealed by immunoblotting with anti-TAP and anti-Strep (five thousand-fold dilution). Expected sizes are 204 kDA for A190-HA and 82 kDa for IN1-strep (WT and mutants). **H.** Genome browser visualization of HA-IN1 occupancy at the *RDN1* locus encoding ribosomal RNA genes. The 100 to 200 *RDN1* repeats of the yeast genome on chromosome XII are aggregated into two repeats. *RDN37-1* and *RDN37-2* are transcribed by RNA Polymerase I as a 35S precursor rRNA. *RDN5-1* and *RDN5-2* are transcribed by RNA Polymerase III to give the 5S RNA. Occupancy of WT HA-IN1 and K_617_A, S_621_A and L_622_A HA-IN1 mutants is represented in each panel. Control, as described for panel D. Values obtained from ChIP seq analysis have been normalized for each condition (WT and mutant IN1) to input and adjusted in log2 RPKM.

To determine if AC40 plays a major role in IN1 recruitment to Pol III-transcribed genes, we developed IN1 chromatin-immunoprecipitation (ChIP) experiments to assay the effect of the K_617_A, S_621_A, and L_622_A mutations on recruitment. WT IN1 and the various mutants were tagged at their N-terminus using a 3xHA epitope tag and expressed from a tetracycline-off promoter. Quantitative PCR revealed significant enrichment of ectopic WT HA-IN1 at all tested Pol III-transcript loci, compared to background level measured on the *GAL1* gene promoter (Figure 3B). In contrast, HA-IN1 mutants that did not interact with AC40 were barely detected at these loci. Thus, recruitment of IN1 to Pol III-transcribed loci depends on its interaction with AC40.

To assess if the genome-wide occupancy of WT IN1 correlates with Ty1 integration site preferences, we performed ChIP sequencing (ChIP-seq) using the same HA-IN1 constructs, with an untagged IN1-expressing strain as control. Analysis of reads mapping to unique sites revealed a strong association of WT HA-IN1 with most nuclear *tDNAs* and the Pol III-transcribed genes *SNR6*, *SNR52*, *SCR1*, *RPR1*, and *RDN5*. Very weak or no HA-IN1 binding was observed for *RNA170* and *ZOD1*, previously shown to have low level of Pol III occupancy (Moqtaderi and Struhl, 2004) (Figure 3C and Table S1). Three *tDNA*s, *tK(CUU)C*, *tM(CAU)C*, and *tD(GUC)N*, were not recovered: These genes are either absent or transcriptionally inactive in the laboratory strain we used (Kumar and Bhargava, 2013; Patterson et al., 2019). HA-IN1 was absent from most Pol II-transcribed genes (Table S1) or present at a much lower level than at Pol III-transcribed genes (Figure 3C). Low HA-IN1 occupancy at Pol II transcribed-genes may be an artifact of ChIP-seq due to the level of expression of these genes (Teytelman et al., 2013). We did not detect significant HA-IN1 binding at other chromosomal loci. The genome-wide distribution of ectopic WT IN1 revealed a strong bias for Pol III-transcribed genes, confirming that the interaction of IN1 with the Pol III is the main driver for targeted integration of Ty1. Under physiological conditions, IN1 is associated with Ty1 cDNA as part of the PIC. In our experimental conditions, IN1 was expressed ectopically at 30°C, a temperature that restricts Ty1 replication. Therefore, our results indicate that the cDNA is not necessary for IN1 recruitment at Pol III. A recent study reached a similar conclusion for the recruitment of Ty3 integrase at *tDNA* genes (Patterson et al., 2019).

ChIP-seq analysis of the HA-IN1 mutants that compromise the IN1-AC40 interaction (K_617_A, S_621_A, and L_622_A) revealed their occupancy was substantially reduced at all Pol III-transcribed genes, as indicated by the lower number of reads corresponding to Pol III-transcribed genes with the three mutants, compare to WT HA-IN1 (Figure 3D), and by a metagene analysis comparing WT and mutant HA-IN1 binding on all the 275 nuclear *tDNAs* (Figure 3E). No similar effect of these mutants was observed at Pol II-transcribed genes (Figure 3F). Quantification by pair-wise Spearman correlation between WT and either K_617_A, S_621_A, or L_622_A IN1 confirmed the apparent stronger decrease in Pol III occupancy of K_617_A HA-IN1 compared to the other mutants (Figures 3E and S2A-B). The metagene analysis indicated a sharp peak around the transcription start site (TSS), which does not coincide with Ty1 integration sites, normally located upstream of Pol III-transcribed genes (Baller et al., 2012; Bridier-Nahmias et al., 2015; Mularoni et al., 2012). This suggests that in addition to the interaction with Pol III, other features may determine Ty1 integration site preference. HA-IN1 occupancy was not completely suppressed in the three mutants, suggesting that the K_617_, S_621_ or L_622_ Ala substitutions may have a residual level of interaction with AC40 not detected by two-hybrid or coIP assay. Alternatively, additional protein-protein interactions, like those previously identified with other Pol III subunits, could contribute to the recruitment of Ty1 integration complex at Pol III transcribed-genes in the absence of the IN1-AC40 interaction (Cheung et al., 2016).

AC40 is common to both Pol I and Pol III, suggesting that IN1 may also interact with Pol I. Consistently, we found that Pol I is associated with IN1 *in vivo*. Pull-down of A190-TAP, the largest subunit of Pol I, retained WT IN1, whereas no association was detected with the IN1 mutants (Figure 3G). The *RDN1* locus is composed of 100-200 tandem repeats of the 35S-precursor *rDNA*, transcribed by Pol I, and the 5S *rDNA*, transcribed by Pol III (Dammann et al., 1993). Analysis of ChIP-seq reads mapping at multiple positions revealed WT HA-IN1 at this locus (Figure 3H). IN1 occupancy may be overestimated, as reads corresponding to all repeats are aggregated on the two copies that are represented in the *S. cerevisiae* reference genome (https://www.yeastgenome.org). However, IN1 mutants compromising the IN1-AC40 interaction —S_621_A, L_622_A, and particularly K_617_A— reduced HA-IN1 occupancy at Pol I-transcribed loci. Thus, IN1, through its interaction with AC40, is also recruited to genes transcribed by Pol I.

Collectively, these results indicate that the IN1-AC40 interaction is necessary for the interaction between IN1 and Pol III and thus its recruitment at Pol III-transcribed genes. IN1 also interacts with Pol I via AC40 and is present at Pol I-transcribed genes.

### AC40 interaction defective Ty1 mutants have altered integration profiles

The reduced association of HA-IN1 mutants with Pol III-transcribed genes did not result in a significant increase in mutant HA-IN1 occupancy at other specific loci, as revealed by ChIP-seq (Table S1). To investigate the integration profile of Ty1 mutants that have an impaired IN1-AC40 interaction, libraries of His^+^ selected *de novo* Ty1 insertion events were generated in cells expressing WT or mutant Ty1*his3*AI elements from the *GAL1* promoter (Barkova et al., 2018). We used an *spt3-101 rad52∆* mutant strain to avoid both trans-complementation of the mutant IN1 by endogenous WT IN1 and Rad52-dependent recombination events. Initially, we performed qualitative PCR to monitor Ty1 insertion events at the *SUF16* tRNA gene and the *SEO1* subtelomeric gene. These genes were identified as hot spots of Ty1 integration in WT and AC40sp loss-of-interaction mutant, respectively (Bridier-Nahmias et al., 2015). In independent cultures expressing WT Ty1*his3*AI, we observed multiple bands upstream of the *SUF16* tRNA gene, characteristic of Ty1-*HIS3* insertion in the three nucleosomes upstream of *tDNA* genes (Bachman et al., 2005; Dakshinamurthy et al., 2010) (Figure 4A). This profile was significantly different for Ty1*his3*AI harboring K_617_A, S_621_A, or L_622_A mutations in IN1, with many fewer integration events upstream of *SUF16*, and increased insertion at *SEO1*, compared to WT Ty1*his3*AI (Figure 4A). This observation suggests that IN1 mutations at K_617_, S_621_, and L_622_ have the same effect on Ty1 integration site targeting as the AC40sp loss-of-interaction mutant.

**Figure 4.**
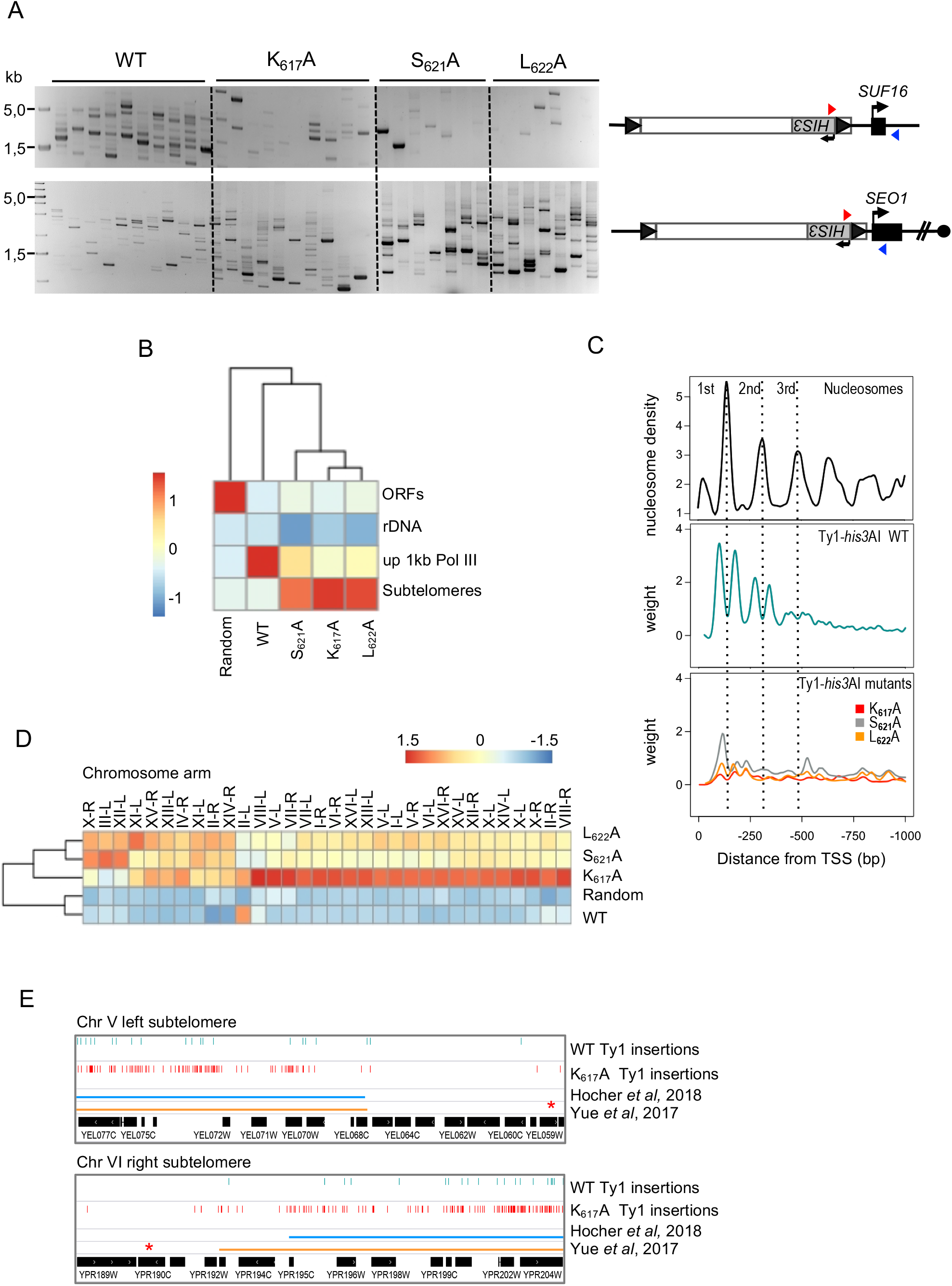
AC40 interaction defective Ty1 mutants have altered integration profiles. **A.** Detection of *de novo* Ty1 insertions upstream of the *SUF16* and *SEO1* genes by PCR using a primer in *HIS3* (red triangle) and a primer in the locus of interest (blue triangle). Ty1 retrotransposition was induced in *spt3-101 rad52Δ* cells transformed with plasmids expressing WT or mutant (IN1 K_617_A, S_621_A and L_622_A) Ty1*his3AI* from the *GAL1* promoter. Total genomic DNA was extracted from His^+^ cells obtained from independent cultures induced for retrotransposition. **B.** Genome-Wide Ty1 insertion frequencies at each genomic feature are clustered in a heatmap. Score is computed in column Z-score. ORFs, all RNA Pol II transcribed genes except gene at subtelomeres; rDNA, one *RDN1* copy; up 1kb Pol III, 1kb upstream of all Pol III transcribed genes; Subtelomeres, genomic coordinates corresponding to chromatin covered by Sir2 and Sir3, when they are co-overexpressed (Hocher et al., 2018); Random, 100.000 random Ty1 computed insertions in the genome. **C.** Ty1 insertion profile upstream of *tDNA*s. Total genomic DNA extracted in (B) was prepared for Ty1 *de novo* integration event sequencing. Ty1 insertions are computed in a 1kb window upstream of all the 275 nuclear *tDNA*s (position 0 in the graph). Each position is divided by the number of insertions at this position (weight). The Smoothing curves indicate the general trend. Nucleosome center positions are from (Brogaard et al., 2012). **D.** Ty1 insertion frequencies for each left and right subtelomere of chromosomes are clustered in a heatmap. Score is computed in row Z-score. Random, as described for panel B. **E.** Genome browser visualization of WT and IN1 K_617_A mutant Ty1-*HIS3* insertions into chromosome V left and chromosome VI right subtelomeres compare to the subtelomere boundaries defined by (Hocher et al., 2018; Yue et al., 2017). Red stars indicate the first essential gene of each subtelomere.

To extend our analysis to the entire genome, we characterized Ty1-*HIS3 de novo* insertion event libraries using high-throughput sequencing. We could discriminate Ty1-*HIS3 de novo* insertions from endogenous elements using six nucleotides in the 3’ LTR that were specific to the Ty1*his3*AI element (Baller et al., 2012). By comparing the Z-score of Ty1 insertions on four non-overlapping features (Figure 4B), we confirmed that WT Ty1 insertions occurred mainly in a 1-kb window upstream of most Pol III-transcribed genes (Table S2), the only exceptions being *tDNAs* that were absent or not transcribed in our strain. There were significantly fewer Ty1-*HIS3* insertions at Pol III-transcribed genes with the three IN1 mutants (Figure 4B). However, the preference for Pol III-transcribed genes was not completely lost, as insertions in these regions were higher than expected if Ty1 targeting was random, supporting the idea that the interaction with Pol III is not fully abolished with these mutants.

WT Ty1-*HIS3* insertions displayed a periodic profile in the region of the three nucleosomes located upstream of Pol III-transcribed genes, with two insertion sites per nucleosome, as seen previously (Figure 4C) (Baller et al., 2012; Bridier-Nahmias et al., 2015; Mularoni et al., 2012). This profile was modified with the three mutants, with the first site of the first nucleosome being less affected with the S_621_A mutant, suggesting that close proximity to the *tDNA* is a determinant of integration. The K_617_A mutant, which had the lowest HA-IN1 occupancy at Pol III-transcribed genes (Figure 3D and 3E), displayed the largest decrease in integration events at Pol III-transcribed genes (Figures 4B and 4C). This correlation suggests that the strength of the IN1-AC40 interaction influences integration efficiency at Pol III-transcribed genes. Concomitant with the decrease in integration at Pol III-transcribed genes, the three IN1 mutants showed an increase in integration events at the end of each chromosome arm, a phenotype that was most pronounced for the K_617_A mutant (Figure 4D). These results are consistent with the redistribution of Ty1 insertions in these regions observed in the AC40sp loss-of-interaction mutant (Bridier-Nahmias et al., 2015). Ty1 *de novo* insertions at the ends of chromosomes were mostly located in regions defined as subtelomeres, based on heterochromatin specificities or loss of synteny between different *Saccharomyces* strains (Hocher et al., 2018; Yue et al., 2017), suggesting that these subtelomeric regions harbor determinants allowing Ty1 targeting (Figure 4E).

Altogether, these results further support a major role for the IN1-AC40 interaction in Ty1 integration targeting at Pol III-transcribed genes. They also confirm that, when this interaction is compromised, Ty1 insertions are not random but principally occur in subtelomeres.

### The IN1 bNLS targets Ty5 integration at Pol III-transcribed genes

To determine whether the IN1 bNLS sequence is sufficient to confer Ty1 integration site preferences, we transferred the sequence into the Ty5 retrotransposon, which preferentially integrates into heterochromatin at yeast silent mating loci (*HMR* and *HML*) and near telomeres (Zou et al., 1996). Ty5 selectivity relies on an interaction between a hexapeptide (TD5, targeting <u>d</u>omain of Ty5) in the C-terminus of IN5 and the heterochromatin protein Sir4 (Gai and Voytas, 1998; Xie et al., 2001). Exchange of TD5 for the IN1 bNLS in IN5, expressed in a two-hybrid vector, resulted in IN5 interacting with AC40, but not with Sir4 (Figure 5A). WT IN5, IN5 lacking TD5 (IN5_∆TD5_), and IN5_∆TD5+bNLS_ harboring the L_622_A mutation in the bNLS sequence, all failed to interact with AC40, demonstrating that the interaction was strictly dependent on the IN1 bNLS sequence.

**Figure 5.**
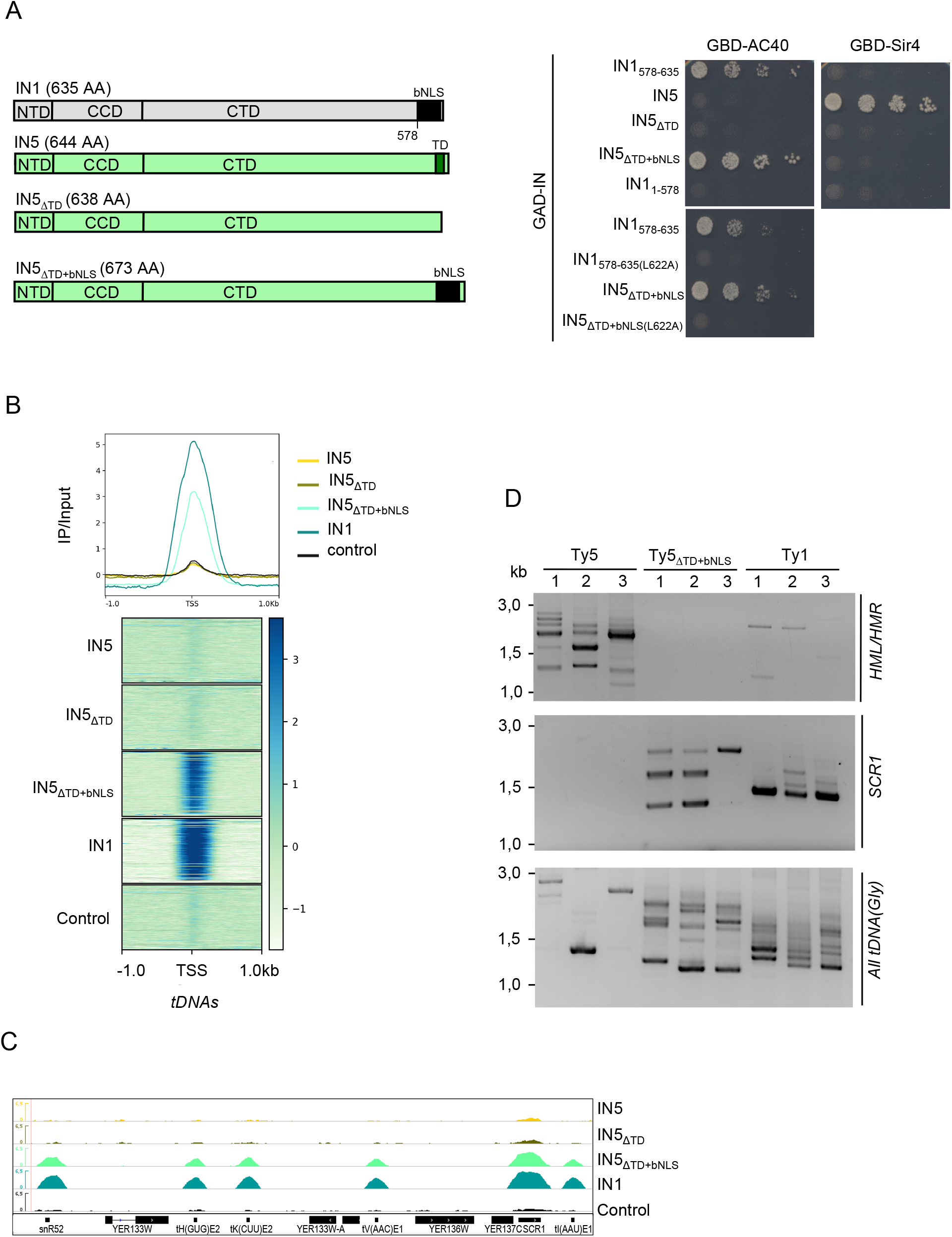
The IN1 bNLS targets Ty5 integration at Pol III-transcribed genes. **A.** Two-hybrid interaction between GBD-Sir4 (left) or GBD-AC40 (right) and different GAD-IN5 or GAD-IN1 constructions. IN1_578-635_ and IN1_1-578_ are used as positive and negative control for interaction with AC40, respectively. IN5 and IN5_ΔTD5_ are used as positive and negative control for interaction with Sir4, respectively. Cells were plated in five-fold serial dilutions on DO-Leu-Trp-His plates to detect interaction. No growth or protein expression defects were detected (Figure S5A-B). **B.** Genome-wide occupancy profiles (top) and heatmaps (bottom) of IN5, WT, and indicated mutants, +/− 1kb upstream and downstream of all *tDNA*s. Average ChIP-seq signals have been computed per 10bp bin, normalized to the input and adjusted in RPKM. **C.** Genome browser visualization of different HA-IN occupancy for chromosome V (chrV:431129..443275). Occupancy of WT IN5, IN5_ΔTD_, IN5_ΔTD+bNLS_ and WT IN1 is represented in each panel. Control, anti-HA immunoprecipitation of chromatin extracts expressing IN1-Strep. The region contains four *tDNAs* (tH(GUG)E2, tK(CUU)E2, tV(ACC)E1, tI(AAU)E1) and two ncRNA genes transcribed by Pol III (*SNR52* and *SCR1*). Values obtained from ChIP seq analysis have been normalized to input and adjusted in RPKM. **D.** Detection by PCR of Ty5, Ty5_ΔTD5+bNLS_ or Ty1 de novo integration events at the *HMR* and *HML* loci, *SCR1* and upstream of all glycine *tDNA*s using a primer in *HIS3* and a primer in the locus of interest. Retrotransposition was induced for three to four days at 20°C in *spt3-101* cells transformed with WT or mutated p*GAL1*-Ty5*his3AI* (Ty5 or Ty5_ΔTD5+bNLS_) and p*GAL1-* Ty1*his3AI* (Ty1). Genomic DNA was extracted from HIS^+^ cells for subsequent PCR assays. Ty5, Ty5_ΔTD5+bNLS_ or Ty1 de novo integration events were detected by using a primer in *HIS3* gene and a primer in the gene of interest. PCR have been performed on genomic DNA extracted from three independent retrotransposition assays.

To establish whether the interaction with AC40 is sufficient to target IN5_∆TD5+bNLS_ to Pol III-transcribed genes, we performed ChIP-seq in strains ectopically expressing HA-tagged IN5, IN5_∆TD5_, and IN5_∆TD5+bNLS_. Metagene analysis of the tagged proteins binding to *tDNA*s revealed a clear enrichment of IN5_∆TD5+bNLS_ at these loci, not detected for IN5 and IN5_∆TD5_ (Figure 5B). IN5_∆TD5+bNLS_ was also present at the other Pol III-transcribed and Pol I-transcribed genes (Figure 5C and S3E). Overall, IN5_∆TD5+bNLS_ genome occupancy profile was very similar to that of IN1 (Pearson correlation of R=0.9 Figure S3F and Table S3), confirming that the IN1 bNLS-AC40 interaction is sufficient for recruitment at Pol III-transcribed genes. We did not detect IN5 enrichment at subtelomeric regions bound by Sir4 (Zill et al., 2010), nor at *HML* and *HMR*, which are Ty5 integration sites, which may be due to the weak association between Ty5 integration sites and Sir4 occupancy (Baller et al., 2011) and reflect a loose and dynamic interaction between IN5 and Sir4 proteins difficult to detect by ChIP.

To explore the impact of the bNLS-TD5 exchange on Ty5 integration site selectivity, we introduced IN5_∆TD5+bNLS_ into a functional Ty5*his3AI* reporter expressed from the *GAL1* promoter. This replacement caused a 10-fold decrease in the frequency of Ty5 retrotransposition but did not inactivate the element (Figure S5G). His^+^ colonies represented *bona fide* integration events, as a similar colony frequency was observed in the absence of homologous recombination (Figure S5G). His^+^ selected *de novo* insertion events were generated for WT and mutant Ty5*his3AI* and the insertion profiles were monitored at specific loci by qualitative PCR (Figure 5D). We used an *spt3-101* strain that does not express endogenous Ty1 elements, to avoid interference between insertions of endogenous Ty1 and the mutant Ty5 at Pol-III transcribed genes. The Ty1*his3AI* profile displayed multiple insertion events at the Pol III-transcribed *SCR1* gene and glycine *tDNAs*, whereas no insertions were seen at *HML* and *HMR*. The WT Ty5 profile displayed multiple insertion events at *HML* and *HMR*, whereas no insertions were recovered at *SCR1* and very few were seen at the glycine *tDNA*s. In contrast, Ty5_∆TD5+bNLS_ did not integrate at *HML* and *HMR*, whereas multiple insertion events were detected at the Pol III-reporter genes. The difference in Ty1 and Ty5_∆TD5+bNLS_ banding patterns at Pol III-transcribed genes is probably due to Ty1, but not Ty5, preferentially integrating into nucleosomes, indicating that the Ty1 preference for nucleosomes is not dependent on interaction with AC40.

Together, these data indicate that the IN1 bNLS sequence is sufficient to direct the integration of the Ty5 retrotransposon upstream of Pol III-transcribed genes.

## DISCUSSION

Here, we show that the Ty1 IN1 bNLS plays a critical role in the interaction with the Ty1 tethering factor AC40. We demonstrate that the interaction with AC40 orchestrates the selection of Ty1 integration sites in the genome and that the IN1 bNLS can function as an independent module that targets the integration of another related retrotransposon upstream of the typical Ty1 Pol III-transcribed target genes.

We provide several lines of evidence supporting a major role for AC40 in recruiting Ty1 to both Pol III and Pol I-transcribed genes. First, IN1 interacts directly with AC40 in the absence of other yeast proteins. Second, mutations in the IN1 bNLS that reduce the interaction with AC40 abolish IN1 association with both Pol III and Pol I transcription complexes. Third, there is a concomitant decrease in IN1 occupancy and Ty1 integration at Pol I and Pol III-transcribed loci when the IN1-AC40 is disrupted. Direct interactions were previously observed *in vitro* between IN1 and the C31, C34, and C53 Pol III-specific subunits (Cheung et al., 2016) but their precise role in the recruitment of the IN1 complex *in vivo* was not determined. The redistribution of Ty1 integration into subtelomeres, seen in the absence of the IN1-AC40 interaction ((Bridier-Nahmias et al., 2015) and this study) was not observed in a *rpc53∆2-280* mutant, which decreases Ty1 integration at the *SUF16* tRNA gene (Cheung et al., 2016). This suggests that either C53 secures IN1 binding to Pol III once IN1 has been recruited by AC40 or helps Ty1 integration at a step downstream of IN1 recruitment. Further studies will be necessary to address the role of the Pol III complex, and especially of C31, C34 and C53, in Ty1 integration.

We show that IN1 is present across Pol I-transcribed genes, and this presence is reduced by mutations that disrupt the IN1-AC40 interaction. Previous genome-wide mapping of Ty1 insertion sites did not reveal a clear pattern of Ty1 insertion into Pol I-transcribed genes at the *RDN1* locus (Baller et al., 2012; Bridier-Nahmias et al., 2015; Mularoni et al., 2012), likely due to the highly repetitive nature of the *rDNA* repeats (Bridier-Nahmias et al., 2015). Ty1 insertion events at these loci seem to be rare (Bryk et al., 1997). This could be due in part to not all the repeats being transcribed by Pol I (Dammann et al., 1993) or to the rapid elimination of *de novo* Ty1 insertions by homologous recombination between *RDN1* repeats. Alternatively, recruitment of IN1 might not be sufficient for Ty1 integration because the process requires additional cofactors or a chromatin structure that is only present at Pol III-transcribed genes.

Our IN1 mutants that disrupt the interaction with AC40 induce the same redistribution of Ty1 insertions at chromosome ends as seen in a AC40sp loss-of-interaction mutant (Bridier-Nahmias et al., 2015). The insertion sites of these Ty1 mutants are scattered throughout subtelomeric regions. Ty5 insertions also occur throughout subtelomeric regions (Baller et al., 2011). This scattered dispersion may explain why we have failed to detect IN1 and IN5 by ChIP-seq in these regions. High-resolution mapping of DNA binding sites will be required to address this point (Hafner et al., 2018; Meers et al., 2019).

To date, the specific retargeting of transposon integration sites has only been observed for Ty1. When the interaction between HIV, MLV, and Ty5 INs and their primary tethering factors (LEDGF/p75, BET proteins, and Sir4, respectively) is altered, the integration of these retroelements at their usual targets decreases substantially, and becomes random (Gai and Voytas, 1998; De Rijck et al., 2013; Schrijvers et al., 2012; Sharma et al., 2013; Wang et al., 2012). Chromosome ends are preferential targets of several TE families in different organisms (Casacuberta, 2017). In *S. cerevisiae*, subtelomeres are devoid of essential genes, are rich in stress responsive genes, and evolve rapidly in response to stress (Snoek et al., 2014) or domestication (Yue et al., 2017). Targeting of Ty1 integration may have evolved to provide a balance between integration into “safe” genomic regions, i.e. *tDNAs*, and integration into fast evolving regions when adaptation is necessary, i.e. subtelomeres. Accordingly, we propose that the IN1-AC40 interaction may be regulated by environmental stress. The observations that Ty5 targeted integration requires phosphorylation of the IN5 targeting domain, which is reduced by stress (Dai et al., 2007), and nutrient starvation regulates the Ty1 replication cycle (Morillon et al., 2000; Todeschini et al., 2005)) both lend support this hypothesis.

The pattern of integration of the Ty1 IN1 mutants largely overlaps in subtelomeric domains (Hocher et al., 2018; Yue et al., 2017). This correlation suggests that specific feature(s) in subtelomeres attract or facilitate Ty1 integration in the absence of the IN1-AC40 interaction. As we failed to detect an interaction between IN1 and Sir4, Ty1 integration into subtelomeres probably involves a mechanism different from that of Ty5. In a *cis*-targeting model, IN1 is tethered to subtelomeres through a weak interaction with a subtelomeric specific co-factor, such that the subtelomeric interaction will only be favored when binding to AC40 is compromised. The dispersed nature of Ty1 insertion sites suggests that the co-factor could be distributed across subtelomeres, like a histone mark specifically enriched in these regions (Hocher et al., 2018). Alternatively, IN1 co-factor could be present at a limited number of subtelomeric sites, and after recruitment, the Ty1 PIC would scan for a chromatin environment favorable for integration. Given Ty1 targets stable nucleosomes upstream of Pol III-transcribed genes (Baller et al., 2012; Mularoni et al., 2012), nucleosome stability could also be a determinant for integration at subtelomeres. A similar two-step targeting model has been proposed to explain the absence of correlation between Sir4 binding sites and Ty5 integration sites in subtelomeres (Baller et al., 2011). In a *trans*-targeting model, the proximity of the subtelomeres with the nuclear pores (Zimmer and Fabre, 2011), through which the Ty1 PIC transits, would facilitate Ty1 integration in these regions, especially when the interaction with AC40 is compromised. Consistent with this hypothesis, mutations in several components of the nuclear pore alter Ty1 integration preferences (Manhas et al., 2018). HIV integration also occurs preferentially in chromatin proximal to the nuclear periphery (Lelek et al., 2015; Marini et al., 2015; Di Primio et al., 2013).

This work demonstrates that the IN1 bNLS functions as an independent module to target integration at Pol III-transcribed genes: The addition of this sequence to Ty5 IN is sufficient to direct Ty5 integration to these loci. Entry of the Ty1 PIC into the nucleus involves the classical import machinery and requires an interaction between importin-α and the IN1 bNLS (McLane et al., 2008). The IN1 NLS consists of two regions of basic amino acids, which are essential for IN1 nuclear import, separated by a linker sequence that also contributes, although to a lesser extent, to import (Kenna et al., 1998; Lange et al., 2011; Moore et al., 1998). The residues that are required for IN1 interaction with AC40 and Ty1 integration upstream of Pol III-transcribed genes cluster in a short peptide within the linker. Mutations in the linker substantially reduce the interaction with AC40 but not Ty1 retrotransposition frequency or IN1 nuclear import (this study). Thus, the IN1 bNLS is involved in nuclear import and integration targeting.

The IN1 linker has been proposed to induce a conformation that facilitates interaction of the two basic amino acid-rich regions with two NLS-binding pockets present in importin-α (Kosugi et al., 2009; Lange et al., 2011). The conformation of the linker when the basic amino acid-rich regions are bound to importin-α could expose the sequence recognized by AC40. Once in the nucleus, interaction with AC40 would assist Ran-GTP to dissociate the IN1-importin-α complex (Passos et al., 2017; Rothenbusch et al., 2012), with nuclear entry and interaction with Pol III being coupled (Figure 6). Such coupling could promote Ty1 targeting to *tDNA*s, as these genes are recruited to the nuclear pore to be transcribed (Chen and Gartenberg, 2014).

**Figure 6.**
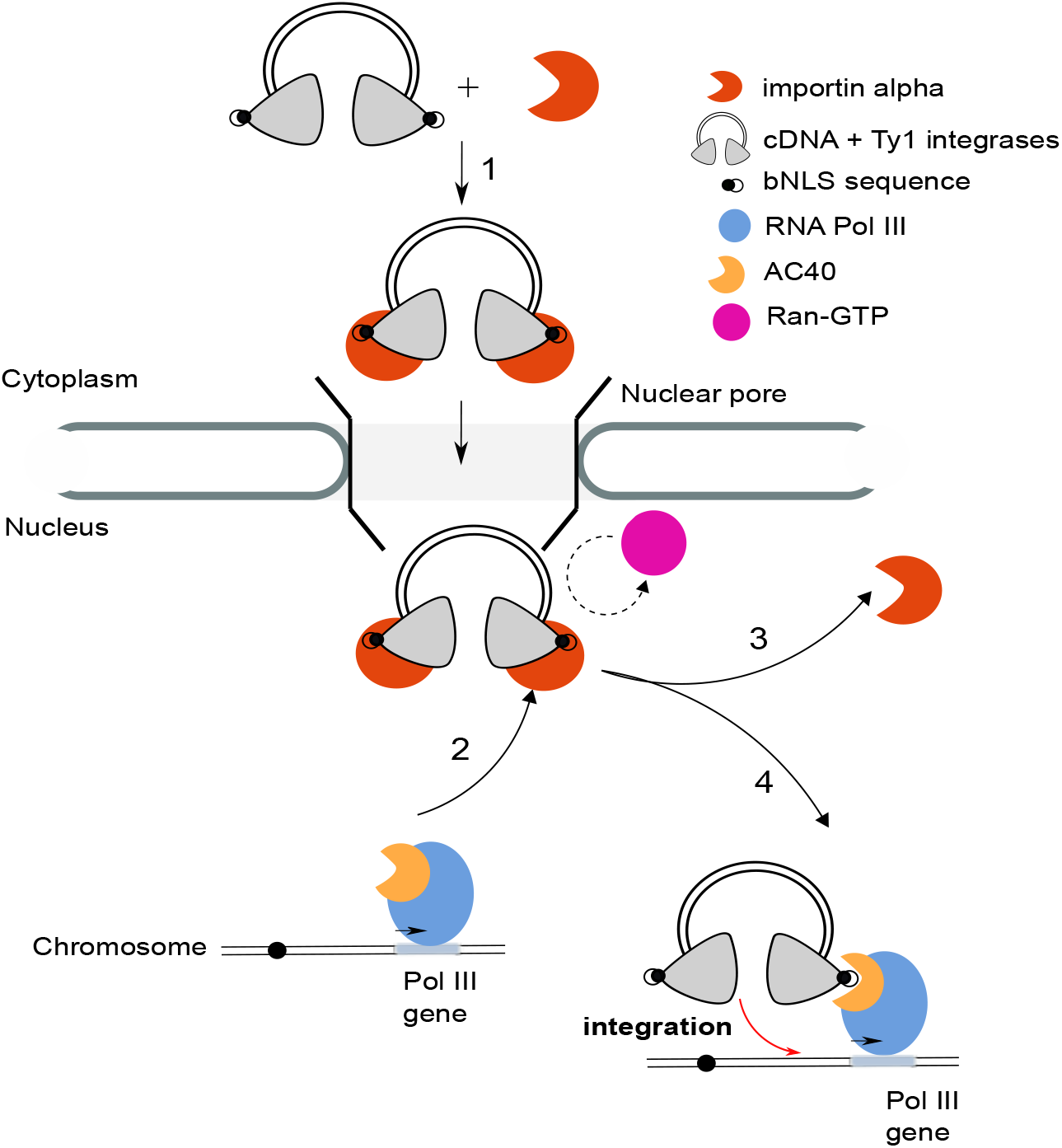
Model coupling the nuclear entry of Ty1 pre-integration complex with Ty1 integration at Pol III-transcribed genes. 1) The Ty1 IN1 bNLS binds to importin-α, allowing the Ty1 pre-integration complex to enter the nucleus. 2) Pol III genes are transcribed at the nuclear pores (Chen and Gartenberg, 2014). The conformation adopted by the IN1 bNLS linker sequence when the two basic amino acid-rich regions are bound to importin-α exposes the sequence recognized by AC40. 3) The binding of AC40 to IN1 assists Ran-GTP dissociate of the IN1-importin-α complex. 4) The association with AC40, in the Pol III complex, targets Ty1 integration upstream of Pol III-transcribed genes.

Retroviral vectors have been used in gene therapy to correct various monogenic disorders. However, these vectors rely on the properties of HIV and MLV integrases, whose preferences for either transcribed genes (i.e. HIV) or promoters (i.e. MLV) make them potentially harmful for the genome (Anguela and High, 2019; Goswami et al., 2019). As Ty1 integration upstream of Pol III-transcribed genes preserves gene integrity (Bolton and Boeke, 2003), and given Pol III transcription and structure, including the presence of AC40, are highly conserved between yeast and humans, adapting Ty1-integration targeting to retroviral integrases could overcome the risk of insertional mutagenesis associated with current MLV and HIV retroviral-based vectors, allowing the development of safer retrovirus-based vectors for use in human gene transfer technologies.

## Acknowledgments

We thank A. Corbett and D. Voytas for plasmids; A. Bridier-Nahmias and members of the laboratory for stimulating discussions; A. Leseur for contribution during her training; E. Fabre, C. Fernandez-Tornero, G. Herrada, B. Palancade, J. Reguera for critical reading of the manuscript. This work was supported by intramural funding from Centre National de la Recherche Scientifique (CNRS), the Université Paris Diderot and the Institut National de la Santé et de la Recherche Médicale (INSERM), and from grants from the Fondation ARC pour la Recherche sur le Cancer (PJA 20151203412), the Agence Nationale de la Recherche through the generic call projects ANR-13-BSV3-0012 and ANR-17-CE11-0025. A.A-L was supported by a post-doctoral fellowship from Fondation pour la Recherche Médicale (FRM-SPF20170938755), A. B. by a post-doctoral fellowship from the ANR through the initiatives d’excellence (Idex ANR-11-IDEX-0005-02) and the Labex “Who am I?” (ANR11-LABX-0071) and I. A. by the PhD program from the CEA. This work has benefited from the facilities and expertise of the high-throughput sequencing platform of I2BC.

## Supplementary Materials

Methods, Supplementary Tables S1-S6, Supplementary Figures S1-S3.

## Author contributions

AAL, AB, CC, CG, HF, IA NP and RM performed the experiments. AAL performed computational analysis. AAL, AB, CC, JA and PL analyzed data and prepared figures AAL, AB, JA and PL wrote the manuscript. JA and PL conceived and supervised the study and secured funding.

