## Supplementary material for "The Ty1 integrase nuclear localization signal is necessary and sufficient for retrotransposon targeting to tRNA genes": Methods, supplemental figures and tables

**Growth media, yeast strains and plasmids construction.** *S. cerevisiae* strains used in this study were grown using standard methods and are listed in Table S4. All plasmids and primers used in this study are reported in Tables S5 and S6. Plasmids were constructed using standard molecular biology procedures (details can be obtained on request). Mutations were introduced in plasmids with Q5® Site directed mutagenesis (NEB). All the constructs were validated by DNA sequencing (Eurofins Genomics).

**Two-hybrid assays.** Two-hybrid assays were performed in strain PJ69-4A which contains the *HIS3* reporter to detect positive interactions (James et al., 1996). Cultures of transformants were grown overnight at 30°C in synthetic complete medium lacking leucine and tryptophan (SC-LEU-TRP) to maintain plasmid selection. Serial dilutions of aliquots of 1 optical density at 600 nm ($OD_{600}$), washed in 1 ml of $H_20$, were plated on SC-LEU-TRP (growth control) or SC-LEU-TRP (interaction), starting from $10^{-1}$ . Plates were incubated 4 days at 20°C. Each two-hybrid assay is a representative example of at least two biological replicates.

**Transposition assays.** To estimate the frequency of retrotransposition in strains transformed by p*GAL1*-Ty1*his3AI*, p*GAL1*-Ty5*his3AI* or their derivatives, four independent transformants of each strain were grown to saturation two days at 30°C in liquid SC-URA containing 2% raffinose. Each culture was diluted thousand-fold in liquid SC-URA containing 2% galactose and grown five days to saturation at 20°C, which is the optimal temperature for Ty1 retrotransposition. Aliquots of cultures were plated on YEPD (100 µl at $10^{-5}$) and SC-HIS (Ty1: 100 µl at $10^{-2}$, Ty5: 1 mL). Plates were incubated for 3 days at 30°C and colonies counted to determine the fraction of [$HIS^+$] prototroph. Retrotransposition frequencies were defined as the mean of at least four experiments, each one performed with four independent transformants.

**Direct fluorescence microscopy.** BY4742 WT cells were transformed with pMET-$GFP_2$-IN1 NLS constructs and pUN100-Nup49-mCherry. Cells were grown overnight in SC-URA-LEU medium at 30°C. Cells were then washed twice in $H_2O$ and resuspended in SC-URA-LEU-MET to induce $GFP_2$-IN1 NLS expression for 6h at 30°C. Live cell imaging was done using a widefield microscopy system featuring a Nikon Ti-E body equipped with the Perfect Focus System and a 100× oil immersion objective. We used an Andor Neo sCMOS camera, which features a large field of view of 276 × 233 µm at a pixel size of 108 nm. We acquired 3D zstacks consisting of 15 frames with z-steps of 300 nm. We used a dual band filter set (eGFP, mRFP) and for each z position acquired two color channels consecutively with an exposure time of 300 ms. The complete imaging system including camera, piezo, LEDs (SpectraX) was controlled by the Andor IQ2 software.

**PCR assays for detection of Ty1 and Ty5 integration events**. After retrotransposition induction as described above, total genomic DNA was extracted from yeast cultures grown at 20°C for 5 days according to classical procedures (Barkova et al., 2018). Double-strand DNA (dsDNA) concentration was determined using Qubit™ fluorometric quantification (Thermofisher Scientific). *De novo* Ty1 insertions upstream of *tG(GCC)C* (*SUF16*) were amplified with PCR primers O-AL27 and O-AB91, and at the *SEO1* subtelomeric gene with PCR primers O-AL27 and O-AL10. *De novo* Ty5 insertions at the *HML* and *HMR* loci, the *SCR1* gene and all glycine *tDNA*s were amplified with O-AL27 combined with O-CG27, O-AL64 or O-AB115 respectively. PCR reaction consisted of 30 ng of dsDNA (or 75 ng for detection of insertions at *SEO1*), 5 µl Buffer 5X, 0,5 µl dNTP 10mM, 0,625 µl of each primer at 20 µM, 0,25 µl of Phusion DNA Polymerase (Thermo Scientific) in a 25 µl final volume. Amplification was performed with the following cycling conditions in ProFlex™ PCR System (Life Technologies) cycler: 98ºC 2 min, 30x [98ºC 10 sec, 60ºC 30 sec, 72ºC 1 min], 72ºC 5 min, hold 4ºC. PCR products were separated on a 1.5% agarose gel.

**Co-immunoprecipitation experiments.** For experiments performed with proteins expressed in *E. coli*, BL21(DE3) Rosetta (Novagen) bacterial cells were transformed with pet28-6xHis-IN1-EPEA (Djender et al., 2014) and pACYC-AC40-Twin-Strep-tag, and streaked on LB agar plates supplemented with ampicillin, chloramphenicol and 1% glucose. Transformants were inoculated in the same medium overnight at 30°C, diluted to 0,05 $OD_{600}$ in fresh 100 ml of LB medium with ampicillin and chloramphenicol without glucose and grown at 30°C to an $OD_{600}$=0.5. Cells were transferred at 24°C for 30 minutes and protein expression was induced by adding 0.5 mM isopropyl 1-thio-β-D-galactopyranoside (IPTG) and growing cells for 3 hours at 24 °C. Cells were harvested at 3500 rpm for 15 min at room temperature, washed with water and resuspended in 1 ml of ice-cold extraction buffer (20 mM Tris-HCl pH 7.5, 300 mM NaCl, 10% glycerol, 0.1% NP40) before adding 1 µg/ml of lysozyme. The extract was kept 10 min on ice and then lysed by sonication (Q700 sonicator, Qsonica) on ice for 5 cycles (output amplitude 10%, 5 sec ON, 40 sec OFF). The lysate was centrifuged at 15 000 rpm at 4°C for 30 min and the supernatant was collected on a separate tube. 800 µl of protein extract was incubated with 40 µl of Magstrep XT beads (Iba-lifescience) for 3 hours at 4°C on a wheel and washed twice with 1 ml of extraction buffer. Bound proteins were eluted with 50 µl of 1X Laemmli sample buffer, separated by SDS-PAGE and analyzed by Western blot using CaptureSelect™ Biotin Anti-C-tag conjugate (Thermofisher) to detect EPEA and anti Strep-Tactin-HRP conjugate (Iba-lifescience). Experiments were reproduced at least 3 times with independent cultures.

Co-IP experiments in yeast cells between Pol III (C160-HA) and IN1-Strep were performed as followed. MW4471 or LV33 yeast strains expressing HA-tagged or untagged C160 respectively were transformed with a 2 micron plasmid expressing WT or $K_{617}$A, $S_{621}$A or $L_{622}$A IN1 mutants from an inducible *GAL1* promoter. Overnight cell cultures were grown to saturation at 30°C in SC-URA to maintain plasmid selection, were diluted at $OD_{600}$=0.001 in 100 mL of SC-URA and containing 2% galactose, and grown at 30°C. At $OD_{600}$=1, cultures were harvested by centrifugation at 4°C. Cell pellets were resuspended in 500 µl of IP extraction buffer (50 mM HEPES-KOH pH 7.5, 300 mM NaCl, 1 mM EDTA, 0.05% NP40, 0.5 mM DTT, 5% glycerol) supplemented with protease inhibitors (ThermoFischer), and lysed with 0.25 mL of glass beads using a vortex (Disruptor Genie®, VWR) for 30 min at 4°C. The whole protein extract (except an aliquot kept for the input control) was incubated with 50 µl of Dynabeads Pan Mouse IgG (Thermo Fischer) coated with anti-HA antibody (12CA5, Roche) 1 hour at 4°C on a wheel. Samples were washed three times with IP extraction buffer. Immunoprecipitated proteins were eluted from beads by boiling samples for 5 min at 95°C with 25 µl of 2X SDS sample buffer and the entire eluted fraction was analyzed by Western blot. Total lanes correspond to 1/20 dilution of the total input engaged in the IP. Proteins were detected with thousand-fold diluted primary antibodies (anti-HA antibody (12CA5, Roche) and anti-strep (Qiagen)) and revealed with ECL (ThermoFischer) and Fusion FX camera (Vilbert-Lourmat). Experiments were reproduced twice with independent cultures.

Co-IP experiments in yeast cells between Pol I (A190-TAP) and ectopically expressed IN1-HBH (HBH is an histidine biotin tag (Tagwerker et al., 2006)) were performed as described for Pol III but with some minor modifications. All the cultures (MGD353-13D or MGD353-13D A190-TAP transformed by pCM185-IN1-HBH) were grown in SC-TRP containing 2% glucose. Immunoprecipitation was performed with Dynabeads Pan Mouse IgG and A190-TAP was detected with 5000-fold diluted primary antibodies (anti-TAP antibody,

Invitrogen). Experiments were reproduced three times with independent cultures.

**Chromatin immunoprecipitation and chromatin immunoprecipitation sequencing.** Yeast strain LV1689 was transformed with pCM185 expressing 3xHA-IN1-EPEA WT or harboring IN1 $K_{617}$A, $S_{621}$A or $L_{622}$A mutations. ChIP was performed as previously described (Harismendy et al., 2003), with minor modifications. Briefly, log-phase cultures (50 ml) in SC-TRP to maintain plasmid selection were cross-linked with 1% formaldehyde for 5 min at room temperature. Cells were pelleted by centrifugation and lysed with glass beads using an orbital shaker (IKA; VRX basic Vibrax; 2200 rpm, 40 min, 4°C). The cell lysate was drawn off the beads and spun for 20 min at 16000 rpm in a microcentrifuge at 4°C. The chromatin pellet was resuspended for 1 hour at 4°C on a rotative wheel and sonicated for 5 cycles (40 sec ON at high level and 20 sec OFF) in a Bioruptor (Diagenode; Denville, NJ, USA). After 1 hour at 4°C on a rotative wheel, the solubilized chromatin was recovered by centrifugation for 15 min at 9 000 rpm at 4°C in a microcentrifuge. The preparation of magnetic beads, immunoprecipitation with anti-HA 12CA5 antibodies, elution from beads and reversal of crosslinking were performed as described (Harismendy et al., 2003). Immunoprecipitated DNA was purified using a QIAquick PCR Purification Kit (Qiagen). The purified DNA samples were analyzed by quantitative real-time PCR using the SYBR® Green PCR master Mix kit and an ABI PRISM 7500 (Applied Biosystems). The results were normalized with the input DNA PCR signals and indicated by relative IP in the graphs. For ChIP-seq analysis, the immunopurified DNA from 3-4 independent experiments were combined after validation by quantitative real-time PCR of *SCR1* and *GAL1* genes. DNA sequencing of 40-nucleotide tags was performed on a GA-IIx, Hi-Seq, or Next-Seq sequencer using the procedures recommended by the manufacturer (Illumina). Input DNA and DNA from ChIP with an untagged strain were used as negative controls. The ChIP-seq data have been deposited to Array Express under accession number E-MTAB-4607.

**ChIP-seq data analysis.** Raw sequence reads quality was controlled with FasqtQC software tool (http://www.bioinformatics.babraham.ac.uk/projects/fastqc/). Reads were trimmed from the adapter sequence and matched to reference genome (SacCer3) using Bowtie2 (Langmead and Salzberg, 2012), with default parameters. Peak calling was performed with MACS2 (Zhang et al., 2008) by comparing ChIP to the corresponding input. Enrichment profile and heatmaps of normalized Reads per Kilo Millions of Mapped reads (RPKM) ratio of IP/input were plotted with Deeptools (Ramírez et al., 2016). Bigwig files were generated for genome browser visualizations. All figures were prepared using R packages (http://www.R-project.org). Graphics were generated using ggplot2 (Wickham, 2016).

**Construction of *de novo* Ty1 insertion libraries for high-throughput sequencing.** LV174 strain (*spt3-101 rad52*Δ) was transformed with pGTy1*his3AI*-SCUF WT (Baller et al., 2012) or pGTy1*his3AI*-SCUF harboring mutations in IN1 sequences (IN1 $K_{617}A$, $S_{621}A$, $L_{622}A$). For each transformant, total genomic DNA was extracted from 70.000 to 100.000 $His^{+}$ colonies recovered from seven to ten independent cultures grown at 20°C in the presence of galactose as described in (Barkova et al., 2018). Fasteris prepared and sequenced Illumina libraries.

**Ty1 Integration data analysis.** Pipeline described in the ChIP-seq Data Analysis section was used for quality check and trimming. To detect *de novo* Ty1 insertions, only reads containing the SCUF sequence and ending with Ty1 LTR sequence were considered. Clean reads were then matched to reference genome (SacCer3) using BWA short aligner (Li and Durbin, 2009) for paired-end reads. PCR duplicates were removed with MarkDuplicates (Picard tools, http://broadinstitute.github.io/picard/) and ambiguous reads ending by Ty1 LTR were discarded. Only properly aligned paired reads (one start and end coordinates) were retained for further analysis. Ty1 *de novo* integrations were assigned to the corresponding genomic features and analyses were performed using in-house R (http://www.R-project.org) pipelines. Graphics were generated using ggplot2 (Wickham, 2016).

#### SUPPLEMENTARY FIGURE LEGENDS

##### Figure S1

**A.** Growth control of cell cultures corresponding to the two-hybrid assay shown in Figure 1B. Two-fold serial dilutions of cell cultures, starting from $10^{-1}$, were plated on DO-Leu-Trp plates to check for growth.

**B.** Validation of the expression of GAD-IN1 fusion proteins for the two-hybrid assay shown in Figure 1B. Whole-cell extract samples were prepared for immunoblotting from 1 $DO_{600}$ of the cell culture by TCA precipitation. GAD-IN1 fusion proteins were detected by Western-blot with anti-HA antibody (12CA5, Roche). All constructs harbour an HA-tag that is present between the GAD and IN1 sequences in the original pACTII vector. Expected sizes of the proteins are 25 kDa for GAD-$IN1_{578-635}$ and GAD-$IN1_{578-635}$ mutants and 82 kDa for GAD-$IN1_{1-578}$.

**C.** Growth control of cell cultures corresponding to the two-hybrid assay shown in Figure 1C. Two-fold serial dilutions of cell cultures, starting from $10^{-1}$, were plated on DO-Leu-Trp plates to check for growth. The complete range of dilutions spotted on DO-Leu-Trp-His is also shown on the right panel.

**D.** Validation of the expression of GAD-IN1 fusion proteins for the two-hybrid assay shown in Figure 1C. Whole-cell extract samples were prepared for immunoblotting from 1 $DO_{600}$ of the cell culture by TCA precipitation. GAD-IN1 proteins were detected by western-blot with anti-GAD antibody (Santa Cruz Biotechnology). Expected sizes of the proteins are 82 kDa for GAD-$IN1_{1-578}$, 27 kDa for GAD-$IN1_{578-635}$ and 24 kDa for GAD-$IN1_{596-630}$.

##### Figure S2

**A.** Genome browser visualization of IN1-HA occupancy for chromosome V (coordinates chrV:431129..443275). Occupancy of WT and mutants HA-IN1, $K_{617}$A, $S_{621}$A and $L_{622}$A is represented in each panel. Control track (black) is an anti-HA immunoprecipitation of chromatin extracts in cells expressing IN1-Strep. The region contains four *tDNAs* (*tH(GUG)E2*, *tK(CUU)E2*, *tV(ACC)E1*, *tI(AAU)E1*) and two ncRNA genes transcribed by Pol III (*snR52* and *SCR1*). Values obtained from ChIP-seq analysis have been normalized to input and adjusted in RPKM.

**B.** Pearson correlation heatmap at *tDNA*s between the following ChIP-seq: IN1 (WT), $K_{617}A$, $S_{621}A$, $L_{622}A$ and control.

**Figure S3**

**A and C.** Growth control of cell cultures corresponding to the two-hybrid assay shown in Figure 5A. Ten-fold serial dilutions of cell cultures, starting from $10^{-1}$, were plated on DO-Leu-Trp plates to check for growth.

**B and D.** Validation of the expression of GAD-IN1 and GAD-IN5 fusion proteins for the two-hybrid assay shown in Figure 5A. Whole-cell extract samples were prepared for immunoblotting from 1 $DO_{600}$ of the cell culture by TCA precipitation. Fusion proteins were detected by Western-blot with anti-GAD antibody (Santa Cruz Biotechnology). Expected sizes of the proteins are 25 kDa for GAD-$IN1_{578\text{-}635}$ and GAD-$IN1_{578\text{-}635}$ mutants and 82 kDa for GAD-$IN1_{1\text{-}578}$, around 100kDa for GAD-IN5 constructions.

**E.** Genome browser visualization of HA-IN1 or HA-IN5 occupancy at the *RDN1* locus encoding ribosomal RNA genes. Control track (black) is an anti-HA immunoprecipitation of chromatin extracts in cells expressing IN1-Strep.Values obtained from ChIP-seq analysis have been normalized for each condition (WT and mutant IN5) to input and adjusted in log2 RPKM.

**F.** Pearson correlation heatmap at *tDNA*s between the following ChIP-seq: IN1, IN5, $IN5_{\Delta TS5}$, $IN5_{\Delta TS5+bNLS}$ and control.

**E.** Retrotransposition frequency, shown on a log-scale of p*GAL1*-Ty5*his3AI* and p*GAL1*-Ty5*his3AI* ($IN_{\Delta TD+bNLS}$) in an *spt3-101* (LV47) or *rad52*Δ strain (LV172). Values are the average of two experiments, each performed with four independent colonies.

### Asif-Laidin *et al.* Figure S1

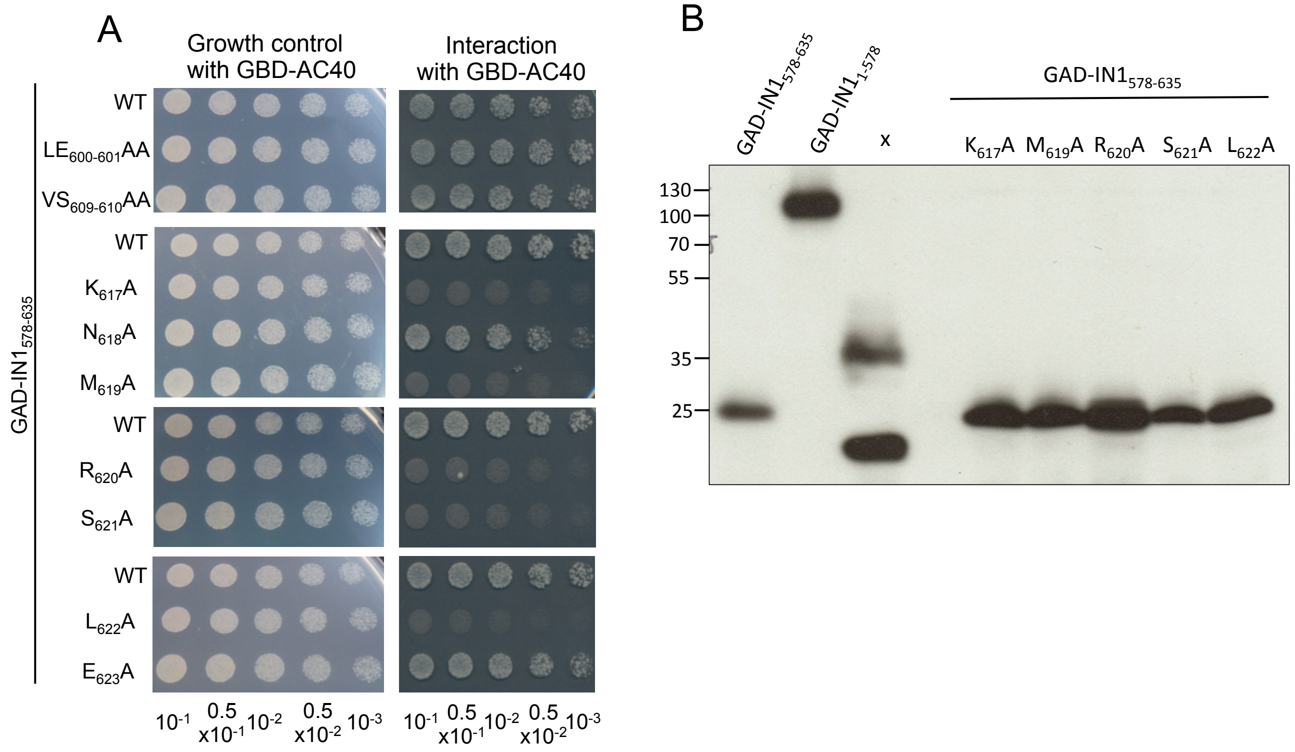


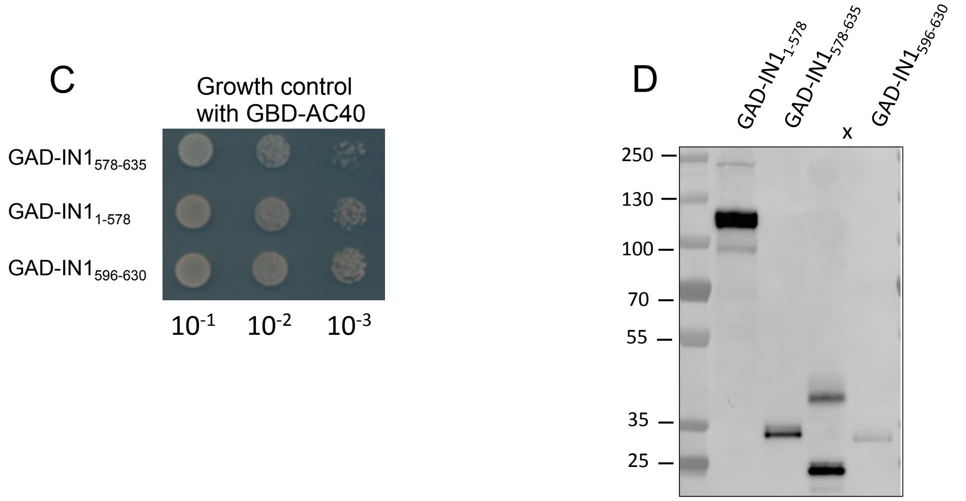

### Asif-Laidin *et al.* Figure S2

A

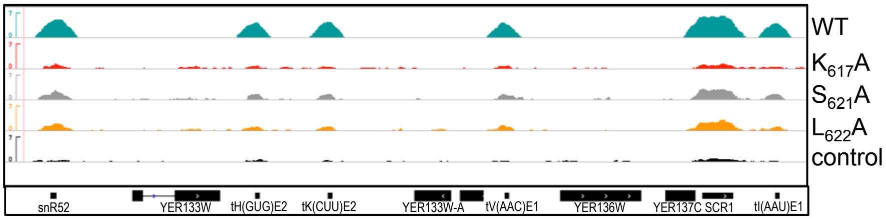


B

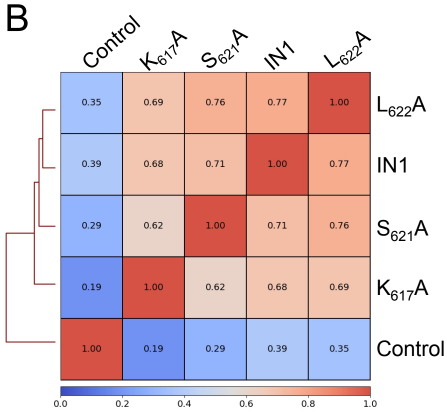

### Asif-Laidin *et al.* Figure S3

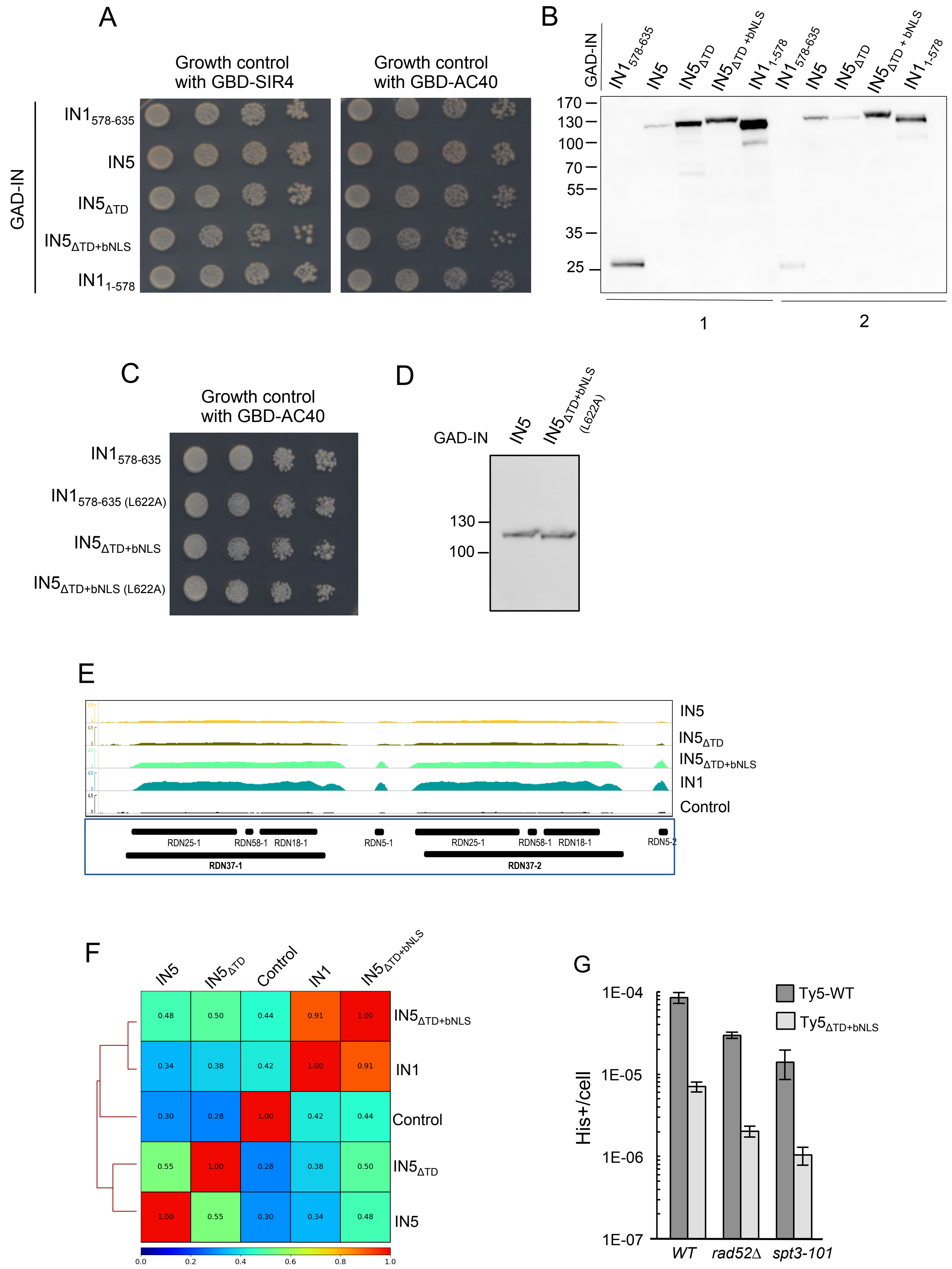

#### Table S4. Strains

| Strain | Genotype | Origin |
|---|---|---|
| BY4742 | *MATα his3Δ1 leu2Δ0 lys2Δ0 ura3Δ0* | Euroscarf |
| LV33 | *MATα ura3Δ851 trp1Δ63 his3Δ200* | (Morillon et al., 2000) |
| LV47 | *MATα ura3Δ851 trp1Δ63 his3Δ200 spt3-101* | (Todeschini et al., 2005) |
| LV172 | *MATα ura3Δ851 trp1Δ63 his3Δ200 Δrad52::TRP1* | This study |
| LV174 | *MATα ura3Δ851 trp1Δ63 his3Δ200 spt3-101 Δrad52::TRP1* | This study |
| LV1689 | *MATa his3Δ1 leu2Δ0 met15Δ0 ura3Δ0 trp1Δ::NatMX* | This study |
| PJ69-4A | *MATα trp1-901 leu2-3,112 ura3-52 his3Δ200 gal4Δ gal80Δ LYS2::GAL1-HIS3 GAL2-ADE2 met2::GAL7-lacZ* | (James et al., 1996) |
| MW4471 | *MATα ura3Δ851 trp1Δ63 his3Δ200 YOR116C-6xHA-KlTRP1* | (Soutourina et al., 2006) |
| MGD353-13D | *MATa ura3-52 trp1-289 leu2-3,112 ade2 arg4* | (Séraphin et al., 1988) |
| MGD353-13D A190-TAP | *MATa ura3-52 trp1-289 leu2-3,112 ade2 arg4 YOR341W-TAP-KanMX* | (Gavin et al., 2002) |

#### Table S5. Plasmids

| Name | Vector | Genotype | Origin |
|---|---|---|---|
| pAT36 | pAS2ΔΔ | 2μ AmpR TRP1 GBD-AC40 | (Bridier-Nahmias et al., 2015) |
| pAT3 | pACTII | 2μ AmpR LEU2 GAD-IN 578-635 | (Bridier-Nahmias et al., 2015) |
| pAT26 | pACTII | 2μ AmpR LEU2 GAD-IN 1-578 | (Bridier-Nahmias et al., 2015) |
| pRM37 | pACTII | 2μ AmpR LEU2 GAD-IN 1-578 Strep | this study |
| pAL4 | pACTII | 2μ AmpR LEU2 GAD-IN 596-630 Strep | this study |
| pAL6 | pACTII | 2μ AmpR LEU2 GAD-IN 578-635 Strep | this study |
| pPL321 | pACTII | 2μ AmpR LEU2 GAD-IN 578-635 LE601-602A | this study |
| pPL322 | pACTII | 2μ AmpR LEU2 GAD-IN 578-635 VS609-610A | this study |
| pPL323 | pACTII | 2μ AmpR LEU2 GAD-IN 578-635 K617A | this study |
| pPL327 | pACTII | 2μ AmpR LEU2 GAD-IN 578-635 N618A | this study |
| pPL328 | pACTII | 2μ AmpR LEU2 GAD-IN 578-635 M619A | this study |
| pPL329 | pACTII | 2μ AmpR LEU2 GAD-IN 578-635 R620A | this study |
| pPL330 | pACTII | 2μ AmpR LEU2 GAD-IN 578-635 S621A | this study |
| pPL331 | pACTII | 2μ AmpR LEU2 GAD-IN 578-635 L622A | this study |
| pPL332 | pACTII | 2μ AmpR LEU2 GAD-IN 578-635 E623A | this study |
| pPL2 | pGAL1Ty1his3AI | 2μ AmpR URA3 pGAL1 Ty1his3AI | (Curcio and Garfinkel, 1991) |
| pCG1 | pGAL1Ty1his3AI | 2μ AmpR URA3 pGAL1 Ty1his3AI IN-K617A | this study |

| | | | |
|---|---|---|---|
| pCG2 | pGAL1Ty1his3AI | 2µ AmpR URA3 pGAL1 Ty1his3AI IN-M619A | this study |
| pCG3 | pGAL1Ty1his3AI | 2µ AmpR URA3 pGAL1 Ty1his3AI IN-R620A | this study |
| pCG4 | pGAL1Ty1his3AI | 2µ AmpR URA3 pGAL1 Ty1his3AI IN-S621A | this study |
| pCG5 | pGAL1Ty1his3AI | 2µ AmpR URA3 pGAL1 Ty1his3AI IN-L622A | this study |
| pRM32 | pGAL1Ty1his3AI | 2µ AmpR URA3 pGAL1 Ty1his3AI IN-D154A | this study |
| pAL9 | pGAL1Ty1his3AI | 2µ AmpR URA3 pGAL1 Ty1his3AI IN-KKR628-630 GGT | this study |
| pPL2-SCUF | pGAL1Ty1his3AI-SCUF | 2µ AmpR URA3 pGAL1 Ty1his3AI-SCUF IN WT | (Baller et al., 2012) |
| pAL12 | pGAL1Ty1his3AI-SCUF | 2µ AmpR URA3 pGAL1 Ty1his3AI-SCUF IN K617A | this study |
| pCG10 | pGAL1Ty1his3AI-SCUF | 2µ AmpR URA3 pGAL1 Ty1his3AI-SCUF IN-S621A | this study |
| pCG11 | pGAL1Ty1his3AI-SCUF | 2µ AmpR URA3 pGAL1 Ty1his3AI-SCUF IN-L622A | this study |
| pAT10 | pGAL1-IN1 | 2µ AmpR pGAL1 IN-strep WT | (Bridier-Nahmias et al., 2015) |
| pCG16 | pGAL1-IN1 | 2µ AmpR pGAL1-IN1-strep K617A | this study |
| pCG20 | pGAL1-IN1 | 2µ AmpR pGAL1-IN1-strep S621A | this study |
| pCG21 | pGAL1-IN1 | 2µ AmpR pGAL1-IN1-strep L622A | this study |
| pCG23 | pACTII | 2µ AmpR LEU2 GAD-IN5 | this study |
| pCG25 | pACTII | 2µ AmpR LEU2 GAD-IN5ΔTS5 | this study |
| pAL1 | pACTII | 2µ AmpR LEU2 GAD-IN5ΔTS5+NLS1 | this study |
| pAL7 | pACTII | 2µ AmpR LEU2 GAD-SIR4 | this study |
| pNK254 | pGAL1Ty5his3AI | 2µ AmpR URA3 pGAL1-Ty5his3AI | (Ke et al., 1997) |
| pAL2 | pGAL1Ty5his3AI | 2µ AmpR URA3 pGAL1-Ty5his3AI ΔTS5+NLS1 | this study |
| pACYC-AC40-strep | pACYC | p15A CmR pT7 AC40-streptagtwin | this study |
| pCM185-IN1-HBH | pCM185 | CEN AmpR TRP1 pTetO7 IN1-HBH | this study |
| pCM185-3HA-IN1-EPEA | pCM185 | CEN AmpR TRP1 pTetO7 3HA-IN1-EPEA | this study |
| pCM185-3HA-IN1K617A-EPEA | pCM185 | CEN AmpR TRP1 pTetO7 3HA-IN1K617A-EPEA | this study |
| pCM185-3HA-IN1S621A-EPEA | pCM185 | CEN AmpR TRP1 pTetO7 3HA-IN1S621A-EPEA | this study |
| pCM185-3HA-IN1L622A-EPEA | pCM185 | CEN AmpR TRP1 pTetO7 3HA-IN1L622A-EPEA | this study |
| pCM185-3HA-IN5 | pCM185 | CEN AmpR TRP1 pTetO7 3HA-IN5 | this study |
| pCM185-3HA-IN5DTD | pCM185 | CEN AmpR TRP1 pTetO7 3HA-IN5DTD | this study |
| pCM185-3HA- | pCM185 | CEN AmpR TRP1 pTetO7 | this study |

| | | | |
|---|---|---|---|
| IN5DTD+NLS1 | | 3HA-IN5DTD+NLS1 | |
| pET28-IN1-EPEA | pET28b | pBR322 KanR pT7 IN1-EPEA | this study |
| pET28-IN1K617A-EPEA | pET28b | pBR322 KanR pT7 IN1K617A-EPEA | this study |
| pET28-IN1S621A-EPEA | pET28b | pBR322 KanR pT7 IN1S621A-EPEA | this study |
| pET28-IN1L622A-EPEA | pET28b | pBR322 KanR pT7 IN1L622A-EPEA | this study |
| pAC1804 | pMET-GFP$_2$-IN NLS | CEN AmpR URA3 pMET-GFP$_2$-IN1 NLS | (McLane et al., 2008) |
| pAC2431 | pMET-GFP$_2$-IN NLS | CEN AmpR URA3 pMET-GFP$_2$-IN1 NLS$_{mut}$ ($_{596}$KKR$_{598}$-AAA and $_{628}$KKR$_{630}$-AAA) | (McLane et al., 2008) |
| pAMA104 | pMET-GFP$_2$-IN NLS | CEN AmpR URA3 pMET-GFP$_2$-IN1K617A NLS | this study |
| pAMA105 | pMET-GFP$_2$-IN NLS | CEN AmpR URA3 pMET-GFP$_2$-IN1S621A NLS | this study |
| pAMA106 | pMET-GFP$_2$-IN NLS | CEN AmpR URA3 pMET-GFP$_2$-IN1L622A NLS | this study |
| pAMA90 | pUN100 | CEN LEU2 AmpR pUN100-Nup49-mCherry | (Chadrin et al., 2010) |

#### Table S6. Primers

| Primer | Sequence 5'- 3' | Gene/Mutation/Plasmid | Reference |
|---|---|---|---|
| SCR1- 4 Forward | GTCCTGGGCAGAGCTGTCT | SCR1-4 | |
| SCR1- 4 Reverse | AAGGTGGAGCCCCTAAGGA | | |
| tRNA Leu Forward | GGTTGTTTGGCCGAGCG | tRNA Leu | |
| tRNA Leu Reverse | TGGTTGCTAAGAGATTCGAACTC | | |
| tRNA Ileu Forward | GCTCGTGTAGCTCAGTG | tRNA Ileu | |
| tRNA Ileu Reverse | TGCTCGAGGTGGGGTTT | | |
| GAL1 Forward | AAAGAAACTTGCACCGGAAA | GAL1 | |
| GAL1 Reverse | GGCCCATATTCGCTTTAACA | | |
| O-RM65 Forward | ACCAAGACACTGGCCTGA | IN1 D154A | |
| O-RM66 Reverse | TGGAGTCGCTGCCCGGCTAAACCG | | |
| O-PL570 Forward | AGATAATGAAACTGAAATTAAGGTATCAC | IN1 LE601-602AA | |
| O-PL571 | GCTGCTGATCTTTTCTTACTGT | | |

| | | | |
|---|---|---|---|
| Reverse | TGATAG | | This Study |
| O-PL572 Forward | TGAAATTAAGGCAGCACGAGACACATG | IN1 VS609-610AA | |
| O-PL573 Reverse | GTTTCATTATCTTCTAATGATCTTTTC | | |
| O-PL574 Forward | GAATATGCGTAGTTTAGAACCTC | IN1 K617A | |
| O-PL575 Reverse | GCAGTATTCCATGTGTCTCGTGATAC | | |
| O-PL580 Forward | CATGCGTAGTTTAGAACCTCCG | IN1 N618A | |
| O-PL581 Reverse | GCCTTAGTATTCCATGTGTCTCG | | |
| O-PL582 Forward | GCGTAGTTTAGAACCTCCGAGATC | IN1 M619A | |
| O-PL583 Reverse | GCATTCTTAGTATTCCATGTGTCTC | | |
| O-PL584 Forward | GAGTTTAGAACCTCCGAGATCG | IN1 R620A | |
| O-PL585 Reverse | GCCATATTCTTAGTATTCCATGTGTC | | |
| O-PL586 Forward | GTTAGAACCTCCGAGATCGAAG | IN1 S621A | |
| O-PL587 Reverse | GCACGCATATTCTTAGTATTCCATGTGTC | | |
| O-PL588 Forward | GGAACCTCCGAGATCGAAGAAAC | IN1 L622A | |
| O-PL589 Reverse | GCACTACGCATATTCTTAGTATTCC | | |
| O-PL590 Forward | GCCTCCGAGATCGAAGAAACGAATTC | IN1 E623A | |
| O-PL591 Reverse | GCTAAACTACGCATATTCTTAGTATTC | | |
| O-RM45 Forward | CAGTTTGAAAAATGAAAAACCCCGCAAGTTC | pGAD-IN1$_{578-635}$-strep | |
| O-RM46 Reverse | CGGATGGCTCCAGTATCTACGATTCATAGATCTCTC | | |
| O-RM79 Forward | CGGATGGCTCCATTAGTTTGACGAGAATTTATCGG | pGAD-IN1$_{1-578}$-strep | |
| O-RM80 Reverse | CAGTTTGAAAAATGAAAAACCCCGCAAGTTC | | |
| O-RM81 Forward | CGGATGGCTCCAGTATCTACGATTCATAGATCTCTC | pGAD-IN1$_{596-630}$-strep | |
| O-RM82 Reverse | CAGTTTGAAAAATAGGGATCCGAATTCGAG | | |
| O-AL42 Forward | AACAATTCACCTGATTGCAGCTG | IN1 KKR$_{628-630}$ GGT | |
| O-AL43 Reverse | CCGCCCGATCTCGGAGGTTCTAAAC | | |
| O-AL4 Forward | CGATGGAGGAGATTCGATTG | pGAD and pGAL1Ty5his3AI$_{\Delta TS5+NLS1}$ | |
| O-AL5 Reverse | AATACCTCATTTAACGCG | | |
| O-AL6 | TTAGAGCCGCGTTAAATGAGG | | |

| | | | |
|---|---|---|---|
| Forward | TATTTCGTTTCTTCGATCTCGG | | |
| O-AL7 Reverse | GAATTCAATCGAATCTCCTCCATCGAGTAAGAAAAGATCATTAGAA | | |
| O-AL27 Forward | GCAAGAGAGATCTCCTACTTTC | HIS3 | |
| O-AL10 Reverse | CGTTGGTGCTGCATATGTCA | SEO1 | |
| O-AL64 Reverse | CGGCCAGAATTCTCAACGTA | SCR1 | |
| O-AB145 Reverse | AAGCCCGGAATCGAACC | All glycine tDNA | |
| O-CG27 Reverse | ACCAGAGAGTGTAACAACAG | HML/HMR | |
| O-AB91 (SNR33OUT) Reverse | TTTTAGAGTGACACCATCGTAC | SUF16 | (Dakshinamurthy et al., 2010) |
